# Gut-associated lymphoid tissue attrition associates with response to anti-α4β7 therapy in ulcerative colitis

**DOI:** 10.1101/2023.01.19.524731

**Authors:** Pablo Canales-Herrerias, Mathieu Uzzan, Akihiro Seki, Rafael S. Czepielewski, Bram Verstockt, Alexandra Livanos, Fiona Raso, Alexandra Dunn, Daniel Dai, Andrew Wang, Zainab Al-taie, Jerome Martin, Huaibin M. Ko, Minami Tokuyama, Michael Tankelevich, Hadar Meringer, Francesca Cossarini, Divya Jha, Azra Krek, John D. Paulsen, M. Zuber Nakadar, Joshua Wong, Emma C. Erlich, Emily J. Onufer, Beth A. Helmink, Keshav Sharma, Adam Rosenstein, Grace Chung, Travis Dawson, Julius Juarez, Vijay Yajnik, Andrea Cerutti, Jeremiah Faith, Mayte Suarez-Farinas, Carmen Argmann, Francesca Petralia, Gwendalyn J. Randolph, Alexandros D. Polydorides, Andrea Reboldi, Jean Frederic Colombel, Saurabh Mehandru

## Abstract

Targeting the α4β7-MAdCAM-1 axis with vedolizumab (VDZ) is a front-line therapeutic paradigm in ulcerative colitis (UC). However, mechanism(s) of action (MOA) of VDZ remain relatively undefined. Here, we examined three distinct cohorts of patients with UC (n=83, n=60, and n=21), to determine the effect of VDZ on the mucosal and peripheral immune system. Transcriptomic studies with protein level validation were used to study drug MOA using conventional and transgenic murine models. We found a significant decrease in colonic and ileal naïve B and T cells and circulating gut-homing plasmablasts (β7^+^) in VDZ-treated patients, pointing to gut-associated lymphoid tissue (GALT) targeting by VDZ. Murine Peyer’s patches (PP) demonstrated a significant loss cellularity associated with reduction in follicular B cells, including a unique population of epithelium-associated B cells, following anti-α4β7 antibody (mAb) administration. Photoconvertible (KikGR) mice unequivocally demonstrated impaired cellular entry into PPs in anti-α4β7 mAb treated mice. In VDZ-treated, but not anti-tumor necrosis factor-treated UC patients, lymphoid aggregate size was significantly reduced in treatment responders compared to non-responders, with an independent validation cohort further confirming these data. GALT targeting represents a novel MOA of α4β7-targeted therapies, with major implications for this therapeutic paradigm in UC, and for the development of new therapeutic strategies.

## Introduction

Vedolizumab (VDZ) is an α4β7-integrin targeting monoclonal IgG1 antibody used in the treatment of ulcerative colitis (UC) and Crohn’s disease (CD)^1, 2^. Among adaptive immune cells, memory T cells and plasma cells demonstrate the highest expression of α4β7, with naïve T and B cells expressing lower frequencies of the integrin^3, 4^. Imprinting of α4β7 on immune cells, initiated within ‘inductive sites’ such as Peyer’s patches (PP) and mesenteric lymph nodes (MLN), is enhanced by the local retinoic acid- and TGF-β-enriched milieu^5^. Further, MAdCAM-1, the ligand for α4β7-integrin is predominantly expressed on intestinal high endothelial venules (HEV)^6^. Accordingly, PP- and MLN-primed T and B cells express α4β7^5, 7^ and preferentially home to the ‘effector sites’ (lamina propria) of the gastrointestinal (GI) tract^8^.

Although VDZ is a frontline drug in the management of UC, our understanding of its mechanism(s) of action (MOA) remains imprecise. The leading hypothesis that VDZ inhibits the migration of pro-inflammatory T cells to the intestinal effector sites remains unsubstantiated by studies^9^, since the relative abundance of lamina propria-resident CD4^+^ and CD8^+^ T cells remains comparable post-treatment to the respective pre-treatment cell frequencies^9^. Further, unlike tumor necrosis factor (TNF) inhibiting drugs, where therapeutic drug monitoring has become a critical component of the treatment paradigms^10^, VDZ levels, anti-VDZ antibodies and α4β7-receptor occupancy focused studies^11^ do not provide clear correlates of therapeutic response to VDZ. Therefore, a better understanding of the MOA of VDZ is critically needed to define biomarkers of drug-response and to inform the development of a number of oral^12^ and parenteral agents targeting the α4β7-MAdCAM-1 therapeutic axis for the management of IBD.

In a small cohort of 6 HIV-1 infected individuals with quiescent IBD, we have previously observed that VDZ targets lymphoid structures^13^. However, the applicability of these observations to HIV-uninfected IBD patients and the underlying mechanism(s) remain undefined. The present report details phenotypic characterization of immune cell perturbations in the peripheral blood and intestinal tissues of UC patients, studied longitudinally as well as cross-sectionally. In particular, we have focused on the effects of anti-α4β7 therapy on intestinal organized lymphoid structures, including isolated lymphoid aggregates and Peyer’s patches, hereafter referred to as gut-associated lymphoid tissues (GALT). We have explored immune cell dynamics in murine studies, using both conventional and transgenic photoactivable models to define intestinal immune cell alterations following anti-α4β7 antibody administration. Finally, we have defined clinical correlates of VDZ-response in two independent cohorts of patients followed longitudinally post-therapy. Altogether, herein we provide a novel appreciation of the role of immune inductive sites in VDZ-treated patients that will inform ongoing and future therapeutic paradigms for IBD.

## Results

### Anti-α4β7 therapy results in a significant decrease of naïve B and T cells in the intestinal mucosa of patients with UC

Patients with UC initiating therapy with VDZ (n=56) were prospectively enrolled, with TNF inhibitor (TNFi)-treated patients (n=10) and untreated UC patients (n=17) serving as controls (COHORT1 characteristics in Table S1). Immune cell composition of ileum and colon was studied using multiparameter flow cytometry in a subset of patients (gating strategy in Figure S1). Total (non-plasma cell) B cells (CD45^+^CD19^+^CD38^-^) were significantly decreased in the colon of VDZ-treated patients but not in TNFi-treated patients (Figure 1A, B). Among B cell subsets, naïve B cells (CD45^+^CD19^+^CD38^-^IgD^+^IgM^+^) were significantly reduced in the ileum and colon of VDZ-treated but not TNFi-treated patients as compared to untreated patients, while switched memory B cells (CD45^+^CD19^+^CD38^-^IgD^-^IgM^-^) were not significantly different between the UC controls, VDZ-treated and TNFi-treated groups (Figure 1A, B). Among patients followed longitudinally with post-VDZ biopsies, there was a significant reduction in naïve B cell frequency in both ileum and colon (Figure 1B, bottom). Notably, 11 out of 12 patients with colonic biopsies had a major drop in the frequency naïve B cells and total B cells after VDZ therapy.

**Figure 1.**
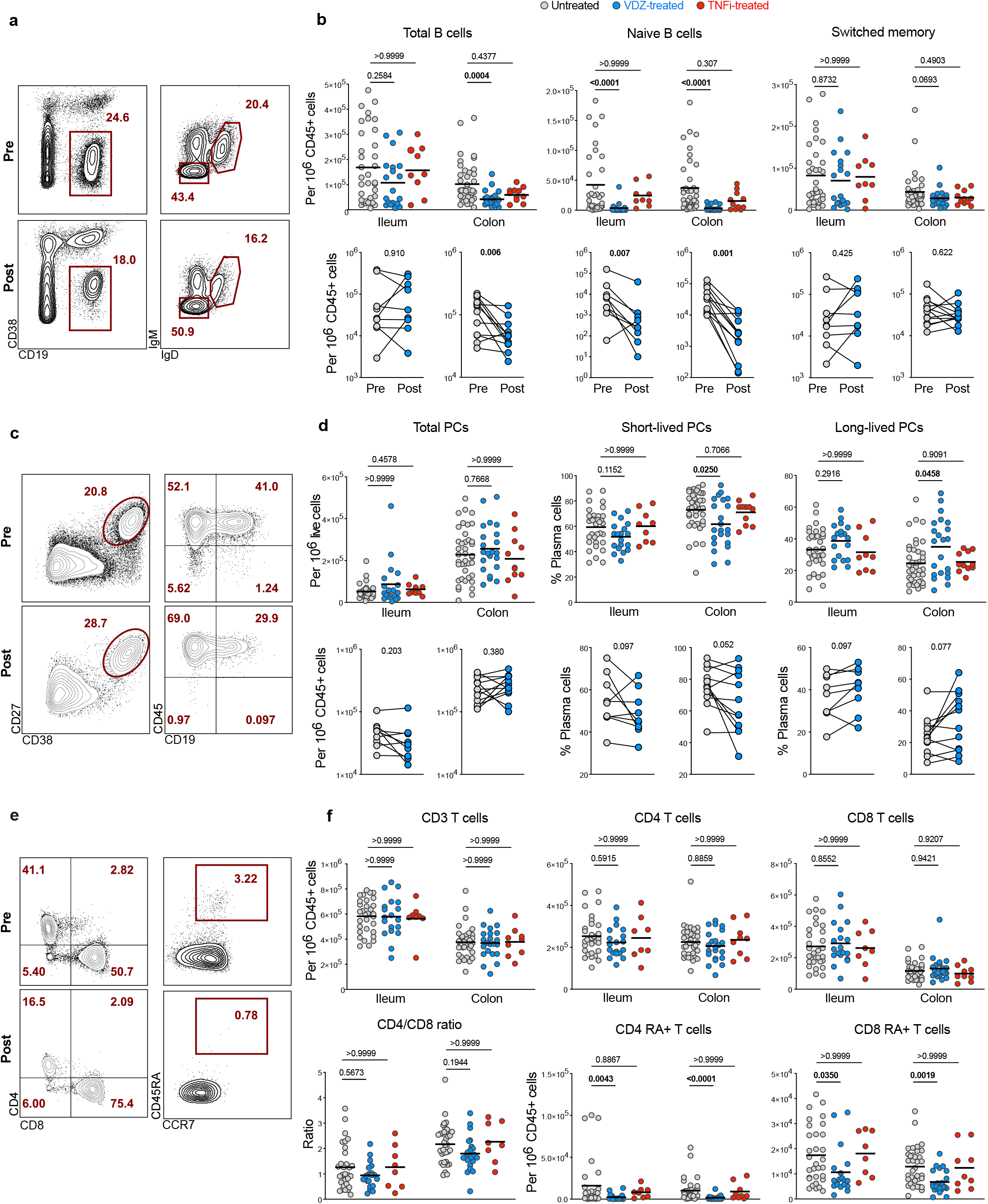
Anti-α4β7 treatment is associated with decreased frequency of naïve B and T cells in the intestines of patients with UC. **a**, Representative flow cytometry (FC) plots for total (non-PC) B cells (CD45^+^CD19^+^CD38^-^), switched memory B cells (CD45^+^CD19^+^CD38^-^IgM^-^IgD^-^) and naïve B cells (CD45^+^CD19^+^CD38^-^IgM^+^IgD^+^) from the ileum of a UC patient pre- and post-VDZ. **b**, Frequency of total B cells, naïve B cells and switched memory B cells from ileum and colon of untreated (n=34), VDZ-treated (n=21) or TNFi-treated UC patients (n=10) (top panels). Longitudinal analysis of paired biopsy samples from UC patients taken pre- and post-VDZ therapy (bottom panels). **c**, Representative FC plots showing total plasma cells (CD27^+^CD38^++^) and their expression of CD45 and CD19 in the left colon of a UC patient pre- and post-VDZ. **d**, Frequency of total plasma cells, short-lived (CD45^+^CD19^+^) and long-lived (CD45^+^CD19^-^) plasma cells in terminal ileum and left colon of UC patients. **e**, Representative FC plots showing the expression of CD4 and CD8 on CD3^+^ T cells (left panel) and the expression of CD45RA on CD4 T cells (right panel) in the ileum of a UC patient pre- and post-VDZ. **f**, Frequency of CD3 T cells, CD4 and CD8 T cells (top panel), as well as naïve (CD45RA^+^) CD4 and CD8 T cells and CD4 to CD8 ratio (bottom panel). Top panels: data are shown as individual data and mean. Non-parametric analysis was done using Kruskal-Wallis test and Dunn’s multiple comparison test, *p* values are indicated. Bottom panels: paired non-parametric analysis was done using Wilcoxon test, *p* values are indicated.

Next, we investigated the frequency of total plasma cells (PC) (CD27^+^CD38^hi^), short-lived PC (CD38^hi^CD27^+^CD45^+^CD19^+^) and long-lived PC (CD38^hi^CD27^+^CD45^+^CD19^-^) as described by Landsverk et al.^14^, in the ileum and colon (Figure 1C). No significant difference was observed in ileal and colonic total PC frequency between VDZ- or TNFi-treated and untreated UC patients (Figure 1D). However, short-lived PC frequency in the colon was significantly decreased after VDZ but not after TNFi therapy (Figure 1D). Furthermore, we observed a non-significant decrease in short-lived PC and non-significant increase in long-lived PC in both the ileum and colon in patients followed longitudinally post-VDZ (Figure 1D).

Among changes in T cells after VDZ, and consistent with previous data^9^, we did not observe significant differences in the frequency of total T cells, CD4^+^ T cells, CD8^+^ T cells and CD4:CD8 T cell ratio in either the ileum or colon of VDZ-treated or TNFi-treated patients compared to untreated UC patients (Figure 1F). However, among T cell subsets, naïve CD4^+^ T cells (CD3^+^CD4^+^CD45RA^+^CCR7^+^) and naïve CD 8^+^ T cells (CD3^+^CD8^+^CD45RA^+^CCR7^+^) were significantly decreased in both ileum and colon in VDZ-treated patients, but not TNFi-treated patients compared to untreated UC patients (Figure 1F). Overall, immunophenotypic examination of the ileum and colon in UC patients treated with VDZ and TNFi demonstrates a VDZ-specific and significant reduction of naïve B and T cells, two cell types that are found predominantly in GALT^15^.

### Anti-α4β7 therapy results in a significant decrease in circulating gut-homing plasmablasts in patients with UC

Next, we studied the evolution of circulating B and T cell subsets, as well as β7-integrin-expressing cells (defined in Figure S2), at week 0 (pre-VDZ) and week 14 post-VDZ initiation, and examined fresh, non-cryopreserved PBMCs by flow cytometry. Notably, and contrary to our expectations, we observed a significant decrease in total plasmablasts (CD10^−^CD19^+/-^ CD27^+^IgD^-^CD38^hi^) after VDZ therapy (Figure 2A-B). Among all plasmablasts, the frequency of gut-homing plasmablasts (CD19^+/-^ CD27^+^CD38^hi^IgD^-^β7^+^) was significantly reduced post-VDZ therapy, while the frequency of non-gut homing (β7^-^) plasmablasts was comparable pre-VDZ and post-VDZ (Figure 2A-B). In contrast to the aforementioned change in plasmablasts, we did not observe changes in total B cells (CD19^+^), switched memory B cells (CD19^+^CD38^-^IgM^-^IgD^-^CD27^+^) or naïve B cells (CD19^+^CD38^-^IgM^+^IgD^+^) in VDZ-treated patients compared to untreated UC controls (Figure 2C-D).

**Figure 2.**
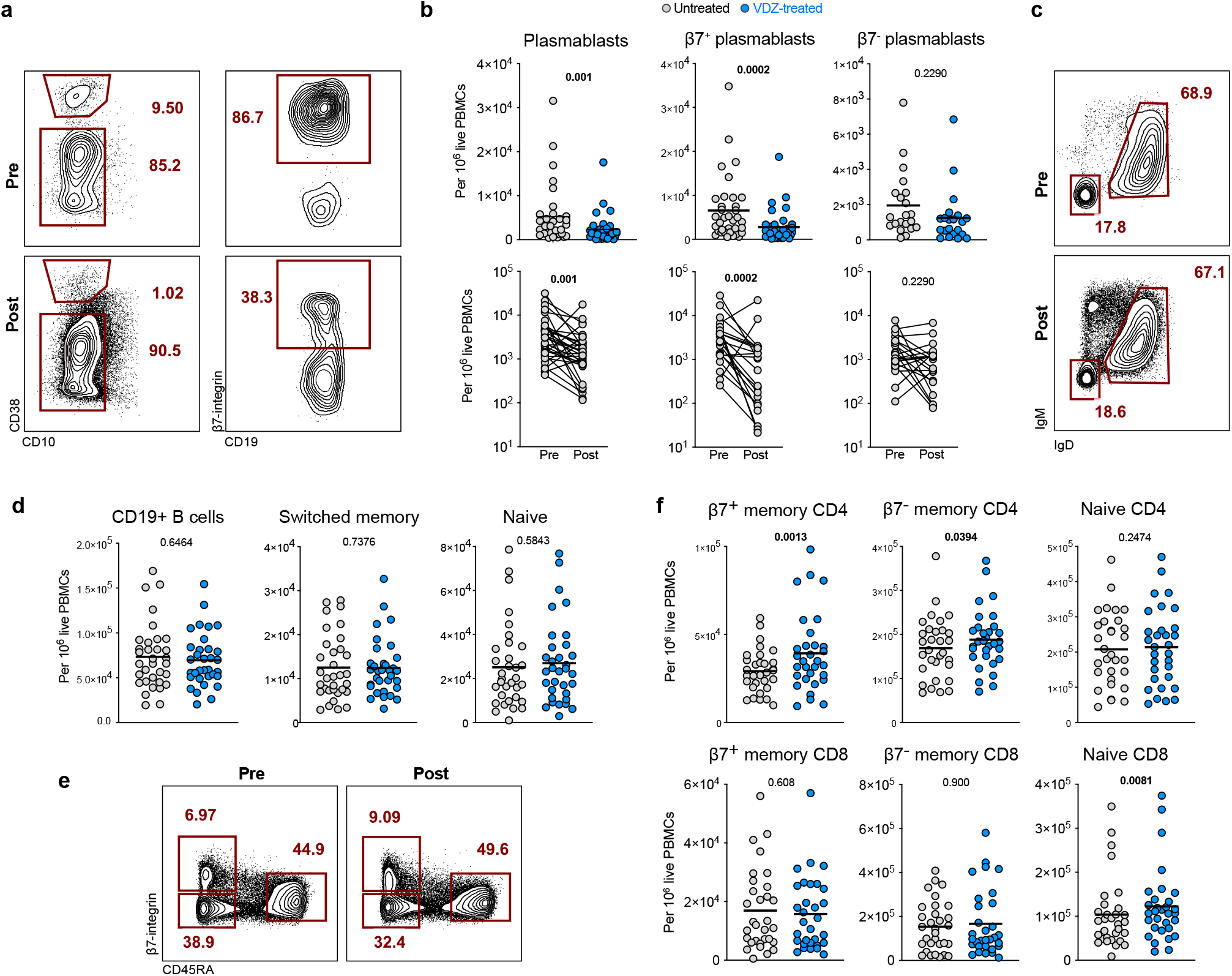
The frequency of gut-homing β7^+^ plasmablasts decreases in peripheral blood of UC patients after VDZ therapy. **a**, Representative FC plots showing the frequency of circulating plasmablasts (CD19^+/int^CD38^hi^CD10^−^) and the percentage of β7^+^ plasmablasts in a patient at week 0 (pre-VDZ) and week 14 (post-VDZ) after initiation of VDZ therapy. **b**, Frequency of total plasmablasts, β7^+^ and β7^-^ plasmablasts in circulation from VDZ-treated UC patients. Longitudinal samples were taken at weeks 0 and 14 after starting VDZ. **c**, Representative flow plots showing the frequency of circulating switched memory B cells (CD19^+^CD38^hi^CD10^−^IgD^-^IgM^-^) and naïve B cells (CD19^+^CD38^hi^CD10^−^IgD^+^IgM^+^) in a UC patient pre-VDZ) and post-VDZ. **d**, Frequency of circulating CD19^+^ B cells, switched memory B cells and naïve B cells between week 0 and week 14 of VDZ therapy. **e**, Representative flow plots showing the frequency of circulating naïve CD4 cells (CD4^+^CD45RA^+^β7^int^), β7^+^ and β7^-^ memory CD4 T cells (CD4^+^CD45RA^-^) in a UC patient pre-VDZ and post-VDZ. **f**, Frequency of circulating memory and naïve T cells in VDZ-treated patients pre-VDZ and post-VDZ. Data is shown as individual values and mean or as paired before-after plots. Paired non-parametric analysis was done using Wilcoxon test and *p* values are indicated. Unpaired analysis was done using Mann-Whitney test, *p* value is indicated.

Next, we analyzed circulating T cells and defined CD4^+^ and CD8^+^ subsets based on CD45RA and β7 expression, as naïve (CD45RA^+^β7^int^), β7^+^ memory (CD45RA^-^β7^+^) and β7^-^ memory cells (CD45RA^-^β7^-^) (Figure 2E). While the frequency of both β7^+^ and β7^-^ memory CD4^+^ cells increased post-VDZ, the effect was more pronounced on the β7^+^ subset (Figure 2F). On the other hand, no difference was noted in the frequency of naïve CD4^+^ T cells in VDZ-treated patients compared to untreated UC controls. In contrast to CD4^+^ T cells, no significant change in memory CD8^+^ cells was noted in VDZ-treated patients, including β7^+^ and β7^-^ memory cells, although naïve CD8^+^ T cells were higher in VDZ-treated patients compared to untreated controls.

Thus, the effect of VDZ on the peripheral blood compartment is cell-specific. While there is an increase in circulating memory CD4^+^ T cells, here we also report for the first time, a significant reduction in circulating β7^+^ plasmablasts after VDZ therapy, a cell population that is primed in intestinal inductive sites.

### Anti-α4β7 antibody therapy results in an attrition of GALT in mice

Based on human data showing a significant reduction in gut-resident naïve B and T cells (enriched in immune inductive sites^16^) and gut-homing β7^+^plasmablasts in circulation (generated in immune inductive sites^16^), we hypothesized that VDZ has a major impact on the intestinal inductive compartment represented by the GALT. To improve our mechanistic understanding, we focused on a murine model of anti-α4β7 antibody treatment (Figure 3A) and examined isolated PPs by flow cytometry. In C57Bl/6 mice, a rapid (within 24 hours) and significant reduction in PP weight and cellularity (Figure 3B) was noted in animals treated with murine anti-α4β7 antibody (DATK32) compared to isotype control- or anti-TNF antibody (MP6-XT22)-treated animals (Figure 3B). Follicular naïve B cells (CD45^+^B220^+^IgD^+^, gating strategy in Figure S3A), the most abundant cell type within PPs, were significantly reduced in mice treated with anti-α4β7 antibody in contrast to isotype control- or anti-TNF-treated mice, while the frequency of germinal-center (GC) B cells (CD45^+^B220^+^IgD^-^GL7^+^FASL^+^) remained unchanged (Figure 3C).

**Figure 3.**
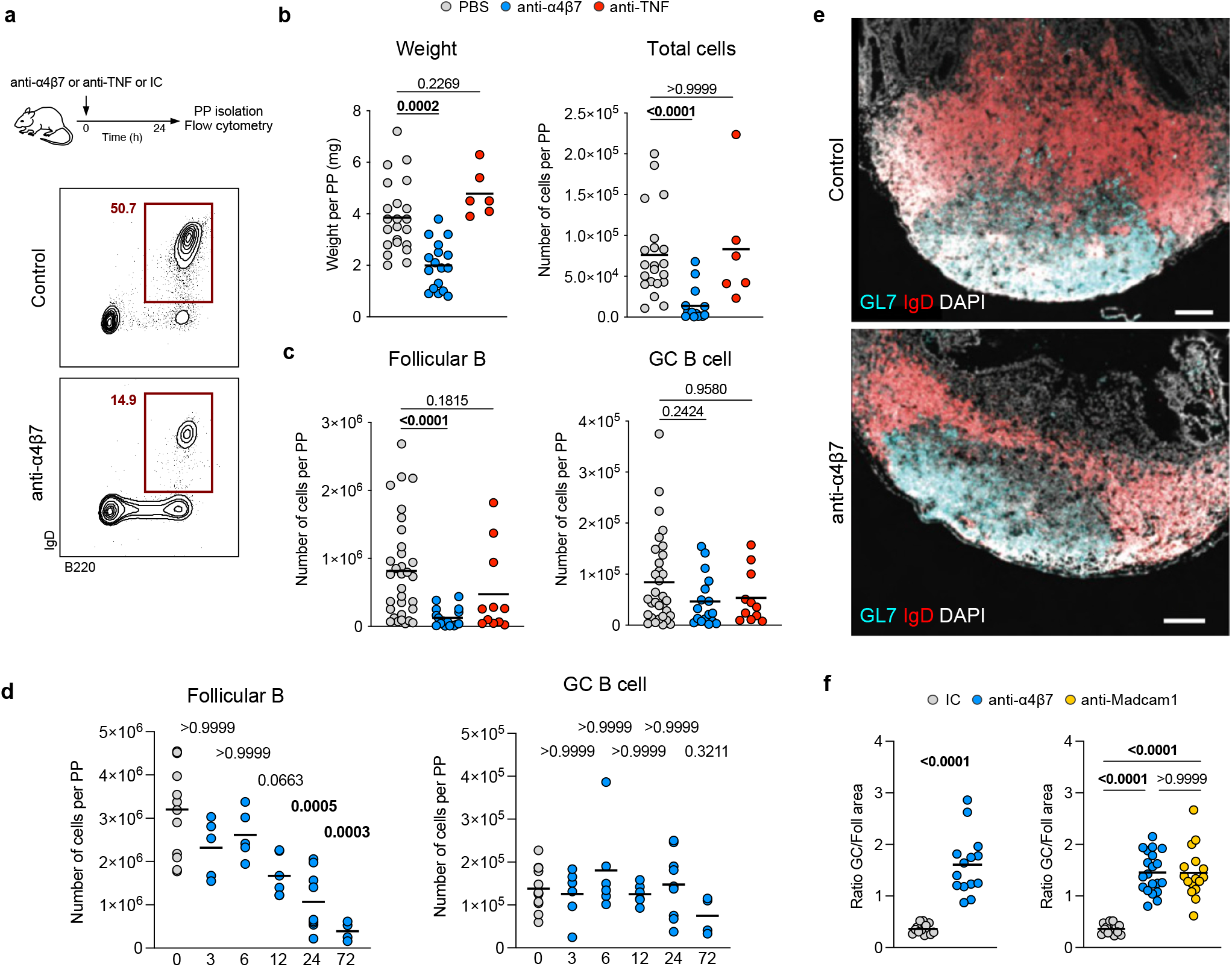
Anti-α4β7 antibody administration results in the attrition of Peyer’s patches in mice. **a**, Mice received one intraperitoneal injection of anti-α4β7 antibody, anti-TNF antibody or PBS antibody, and were sacrificed 24 hours later. Cells from PPs were analyzed by flow cytometry to quantify follicular B cells (CD45^+^B220^+^IgD^+^) and germinal center B cells (CD45^+^B220^+^IgD^-^GL7^+^FASL^+^). **b**, Weight and total cell counts from individual PPs taken 24h after injection. **c**, Frequency of follicular B cells and germinal center B cells from individual PPs. **d**, Frequency of follicular B cells and germinal center B cells from PPs of untreated mice (grey) and at 3, 6, 12, 24 and 72 hours after injection. **e**, Representative immunofluorescence images from PPs stained for IgD (red), GL7 (cyan) and DAPI (grey), from mice treated with anti-α4β7 antibody or Isotype control. Scale bar indicates 100µm. **f**, Ratio of germinal center area (GL7^+^) and follicular area (IgD^+^) from mice treated with anti-α4β7 (blue), anti-MAdCAM-1 (yellow) or isotype control (gray) antibody. Data shown as individual values and mean. Unpaired non-parametric analysis was done using Kruskal-Wallis test and Dunn’s multiple comparisons test. The *p* values are indicated.

Next, we defined the kinetics of cellular change by examining PPs at several timepoints from 3 to 72 hours after anti-α4β7 antibody injection. We observed a progressive loss of follicular B cells, which became significant at 24h after injection (Figure 3D), while the frequency of GC B cells remained unchanged. Analogously, anti-α4β7 administration led to a rapid reduction in PP-resident CD4^+^ and CD8^+^ T cells (Figure S3B). Among T cell subsets, naïve CD4^+^ and CD8^+^ T cells were depleted, with naïve CD8^+^ T cell reduction becoming significant at 24h when compared to baseline (Figure S3C). Finally, we examined the frequency of T follicular helper, PDPN^+^, CD103^+^ and CD11b^+^ cells and found no significant changes as a result of anti-α4β7 administration in the 72h time window (Figure S3D).

We next examined changes in the architecture of PPs by staining for IgD and GL7 to identify the follicular and GC areas respectively. Anti-α4β7 antibody administration resulted in a specific loss of follicular B cell area, while GC area remained unchanged (Figure 3E), resulting in a significant increase in the ratio of GC:follicular area in anti-α4β7-treated mice compared to controls (Figure 3F, left). To establish if the follicular B cell depletion was dependent on α4β7-MAdCAM-1 interaction, we compared mice treated with anti-α4β7 or with anti-MAdCAM-1 antibody to control mice, and observed a significant increase in GC:follicular area ratio in both groups of mice as compared to control mice (Figure 3F). Taken together, this data suggests that anti-α4β7 therapy has a major impact on intestinal lymphoid structures, driven by a loss of follicular naïve B cells and T cells, and attrition of the follicular B cell zone. This process was dependent on blocking the interaction between α4β7 and its ligand MAdCAM-1.

### Anti-α4β7 therapy reduces antigen-specific immune responses in the intestines

To uncover the transcriptional changes in response to anti-α4β7 therapy, we performed single-cell RNAseq on murine PP after one dose of anti-α4β7 antibody or PBS. 15 distinct cellular clusters were annotated on the basis of canonical markers and represented by uniform manifold approximation and projection (UMAP) (Figure 4A). Most cells were mapped to clusters of B cells (Cd79a, Ms4a1, Cd19), plasma cells (Jchain, Mzb1) and T cells (Cd3d, Ms4a4b) (Figure 4B). Additionally, we identified epithelial cells (Epcam), innate lymphoid cells (ILCs) (Kit, Rorc), stromal cells (Col4a1, Ctsl), and dendritic cells (Anxa3, Flt3, Bst2). We next subclassified the B cell compartment into follicular B cells (Ighd, Ighm) and GC B cells (Aicda, Bcl6, Mzb1) (Figure 4C). In PBS-treated mice, follicular B cells represented more than 60% of total cells (Figure 4D). In contrast, there was a major loss of follicular B cells after anti-α4β7 therapy, consistent with our previous data (Figure 3B). All three clusters of follicular B cells were significantly reduced in anti-α4β7 treated mice (Figure 4E), in contrast to other cell clusters (Figure S4).

**Figure 4.**
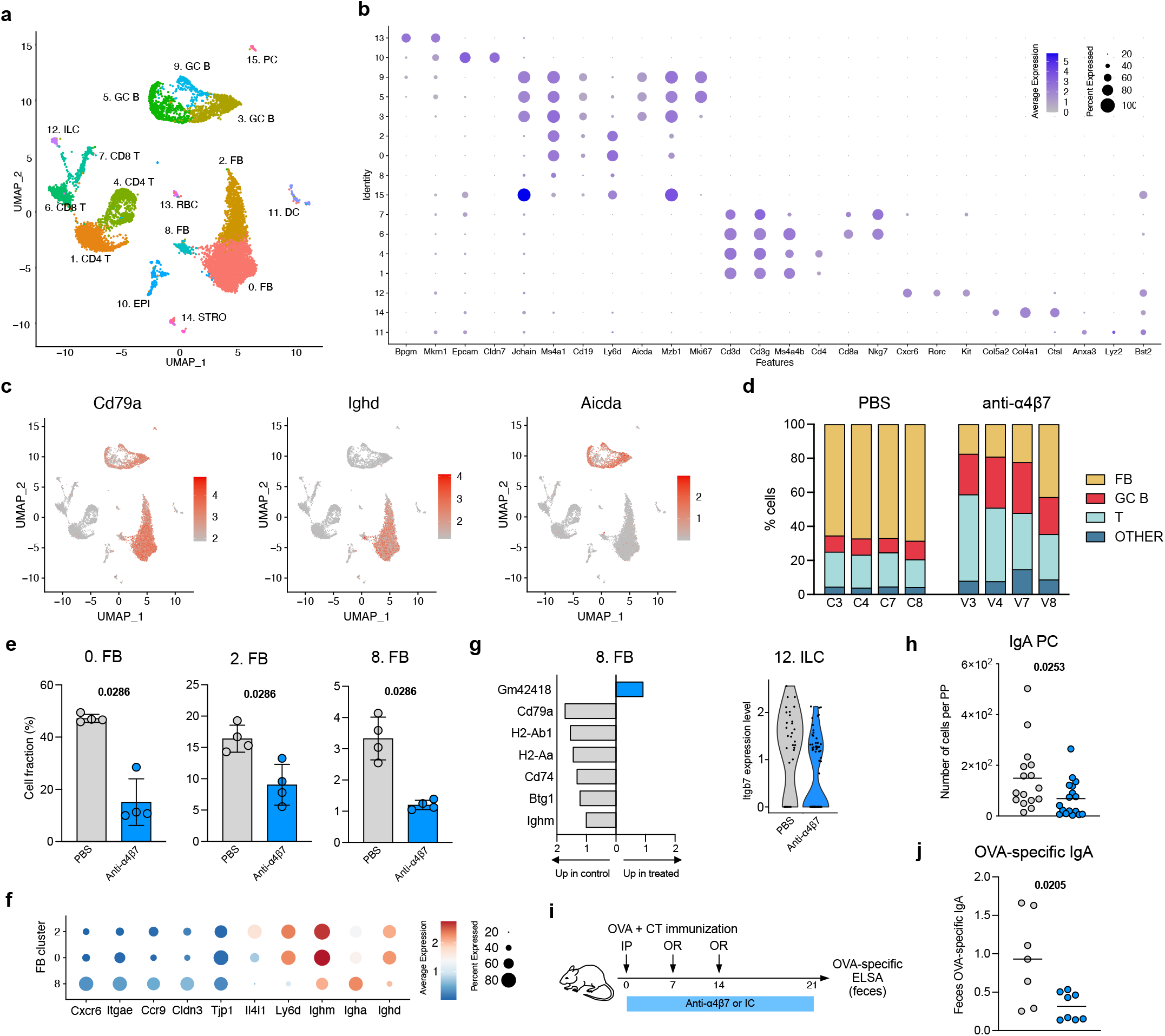
Transcriptional profiling of mouse PPs after anti-α4β7 therapy. **a**, UMAP representation of scRNA-seq analysis of cells isolated from PPs of mice, 24h after anti-α4β7 antibody or PBS administration. **b**, Dotplot showing mean expression and proportion of expressing cells for the indicated genes in each cluster. **c**, UMAP plots showing expression level of the indicated genes. **d**, Bar plots indicating cell composition of PPs from anti-α4β7- and PBS-treated mice. Cells were grouped into follicular B cells (FB), germinal center B cells (GC B), T cells (T) and Others. **e**, Bar plots for the frequency of FB cell clusters among total cells. Data shown as individual values and mean. Unpaired analysis was done using Mann-Whitney test, *p* value is indicated. **f**, Dotplot showing mean expression and proportion of expressing cells for the indicated genes in clusters 0, 2 and 8. **g**, (left) Bar plot of differentially expressed genes in cluster 8 cells between anti-α4β7- and PBS-treated mice. (right) Expression level of Itgb7 in cells from cluster 12. **h**, Frequency of IgA^+^ plasma cells isolated from PPs. **i**, Mice were primed intraperitoneally with OVA + CT (day 0) and boosted orally with OVA + CT (days 7 and 14) and treated with anti-α4β7 or isotype control antibody every 3 days. Feces were collected at day 21. **j**, OVA-specific IgA measured by ELISA. Data shown as individual values and mean. Unpaired analysis was done using Mann-Whitney test, *p* value is indicated.

Next, we sought to better define the changes in the follicular B cell compartment. Cells in clusters 0 and 2 had a profile consistent with follicular naïve B cells^17^, with high expression of Ighd and Ighm, as well as Ly6d and Il4i1 (Figure 4F). Interestingly, cells in cluster 8 had reduced but detectable expression of Ighd and Ighm as well as increased expression of Igha, suggesting that a fraction of these cells had undergone IgA class-switch recombination. Transcriptional profile of cluster 8 cells included genes involved in homing towards the epithelial compartment (Cxcr6, Ccr9) as well as epithelial cell-cell interaction (Itgae, Cldn3, Tjp1). These observations suggest cluster 8 B cells to be associated with the follicle-associated epithelium, possibly in the region of the sub-epithelial dome^18, 19^. We next carried out differential gene expression analysis between anti-α4β7 and PBS-treated mice. In cluster 8 B cells, we found differentially expressed genes (DEGs) involved in MHC class II antigen presentation (H2-Ab1, H2-Aa, Cd74) to be upregulated in control mice compared to anti-α4β7-treated mice (Figure 4G). Among DEGs from other clusters, we found that the β7 gene (Itgb7) was upregulated in controls as compared to treated mice in ILCs (Cluster 12) (Figure 4G), which can also undergo α4β7-dependent homing to gut tissues^20^.

A major function of PP is immune cell priming and induction of antigen-specific immune responses^21^ including IgA^22, 23^. To examine for functional consequences of attrition of lymphoid structures due to anti-α4β7 antibodies, we examined the intestinal IgA response in mice treated with anti-α4β7 antibody. The number of IgA^+^ PC (CD45^+^B220^-^IgA^+^) was significantly reduced 24h after anti-α4β7 antibody administration (Figure 4H). To evaluate the induction of antigen-specific IgA, mice were primed intraperitoneally with 500µg of ovalbumin (OVA) and 0.1µg of cholera toxin (CT) and boosted with 10mg OVA and 10µg CT administrated orally (Figure 4I). OVA-specific IgA antibody was significantly reduced in the stool of anti-α4β7-treated mice compared to control mice (Figure 4J). These findings suggest that anti-α4β7 treatment reduces antigen-specific IgA responses in the intestine, in part by impairing follicular B cell function in PPs.

### Anti-α4β7 antibody blocks B and T cell entry into Peyer’s patches

Next, we asked whether reduced frequencies of PP-resident cells resulted from impaired cellular entry or accelerated egress from the PP. To characterize this process dynamically, we used transgenic mice expressing the photoconvertible protein KikGR(24, 25). Individual PPs (n=3 per mouse) were photoconverted from green to red fluorescence (methods) (Figure 5A). Immediately after photoconversion, most cells contained in photoconverted PPs are K^RED^, while all cells in non-photoconverted PPs remain K^GREEN^. Photoconverted (K^RED^) and non-photoconverted (K^GREEN^) PP-isolated cells (follicular B cells, GC B cells, T cells) were compared between anti-α4β7 antibody- or isotype control-treated mice 20 hours after photoconversion using flow cytometry (Figure 5B).

**Figure 5.**
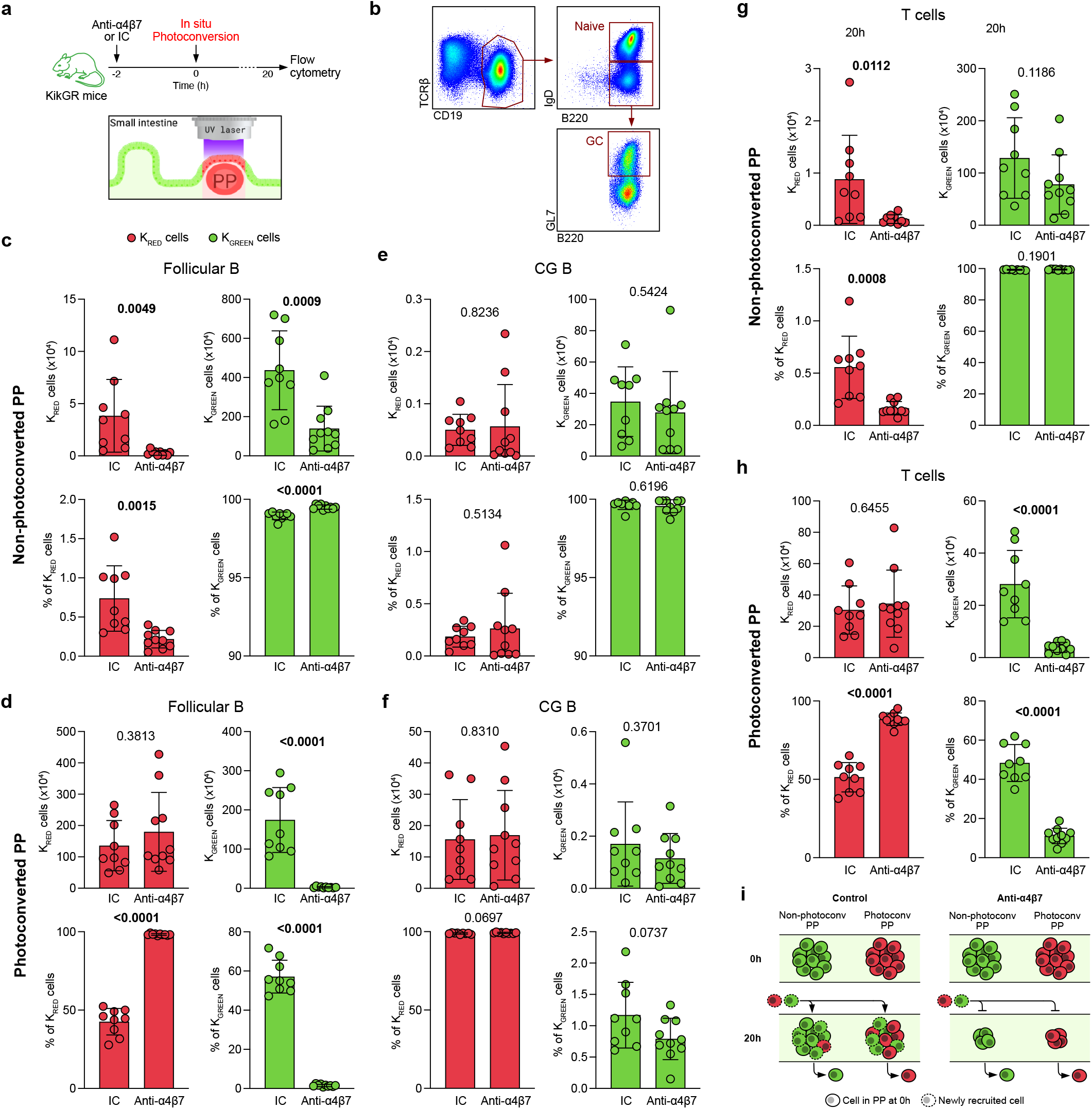
Dynamic monitoring of intestinal lymphocyte trafficking through photoconversion. **a**, Mice were injected with anti-α4β7 or isotype control antibody 2h before photoconversion of 3 PPs using laser light. Organs were harvested 20h later and cells analyzed by FC. **b**, Gating strategy for follicular B cells (CD19^+^B220^+^IgD^+^) and germinal center B cells (CD19^+^B220^+^GL7^+^) from a representative mouse. **c-f**, Absolute cell number and cell frequency of K^RED^ and K^GREEN^ cells out of Follicular B cells (**c, d**) and GC B cells (**e, f**). Cells were isolated from non-photoconverted (**c, e**) and photoconverted (**d, f**) PPs. **g**-**h**, Absolute cell number and cell frequency of K^RED^ and K^GREEN^ T cells from non-photoconverted (**g**) and photoconverted (**h**) PPs. **i**, Model scheme: anti-α4β7 prevents entry of new (red and green) cells into PPs, but does not affect cell egress. This process renders PPs smaller. Data shown as individual values, mean and SD. Unpaired analysis was done using Mann-Whitney test, *p* value is indicated

In non-photoconverted PPs, we observed a significant decrease in the total number of follicular B cells, including both K^RED^ and K^GREEN^ cells, in anti-α4β7-treated mice compared to control mice (Figure 5C, top), in line with our previous data (Figures 3, 4). Interestingly, the proportion of K^RED^ FB cells was significantly reduced after anti-α4β7 treatment, while the proportion of K^GREEN^ cells was significantly greater in anti-α4β7 treated compared to control mice (Figure 5C, bottom). This data suggests that the recruitment of new cells into PPs, including cells coming from photoconverted (red) PPs, was impaired by anti-α4β7 treatment. In photoconverted PPs, similar numbers of K^RED^ follicular B cells were seen in anti-α4β7-treated and control mice (Figure 5D, top), showing that cell egress from PPs was not impacted by anti-α4β7 treatment. In contrast, there was a highly significant reduction in K^GREEN^ follicular B cells in anti-α4β7-treated mice compared to control mice (Figure 5D, top). Furthermore, the Follicular B cell compartment in photoconverted PPs remained almost entirely composed of K^RED^ cells (98%) after anti-α4β7 therapy, while in control mice, more than 50% of FB cells were K^GREEN^ (Figure 5D bottom). These data demonstrate that entry of follicular B cells into PPs is acutely impaired by anti-α4β7 therapy. Our results also serve to highlight the rapid turnover of FB cells in PPs. When we examined the mesenteric lymph nodes, we did not observe any significant differences in FB cell frequencies after anti-α4β7 antibody administration (Figure S5), suggesting that this effect is specific to GALT.

In contrast to follicular B cells, the total number and frequency of GC B cells was comparable between anti-α4β7 treated and control mice in both non-photoconverted (Figure 5E) and photoconverted PPs (Figure 5F), in consistence with our previous data. Analogous to the reduction of FB cells, we observed a significant decrease in K^RED^ total T cells in the non-photoconverted PP of anti-α4β7 treated mice compared to controls (Figure 5G). Additionally, the photoconverted PPs of anti-α4β7 treated mice contained significantly less K^GREEN^ T cells than controls (Figure 5H, top). Finally, the T cell population in photoconverted PPs was dominated by K^RED^ cells (90%) after anti-α4β7 therapy, while control mice had 50% K^RED^ T cells (Figure 5H, bottom). Altogether, these data demonstrate that anti-α4β7 therapy negatively impacts on B and T cell entry into PPs, without affecting cell egress (Figure 5I). Therefore, by altering the balance between cell entry/egress, anti-α4β7 treatment drives a cell deficit in PPs, which progressively promotes attenuation of these structures.

### Attrition of GALT by VDZ is associated with treatment response in patients with UC

Finally, we wanted to study the impact of VDZ on GALT in patients with UC and determine the clinical implications of these findings. We analyzed tissue histology data from a new cohort of UC patients (n=34) where ileal and colonic biopsies were taken pre- and post-VDZ therapy, and selected tissue sections where lymphoid aggregates could be identified (COHORT 2 characteristics in Table S2). TNFi-treated patients were used as controls (n=26). We first determined the mean surface area of all identified lymphoid aggregates per patient and found that there was a trend towards a decrease in the overall size of lymphoid aggregates post-VDZ (Figure 6A-B), while the number of lymphoid aggregates per biopsy remained unchanged (Figure 6B). To determine if lymphoid aggregate size reduction was different between treatment responders and non-responders (response defined as lack of histological inflammation), we compared the mean size of lymphoid aggregates between treatment responders and non-responders (Figure 6A, C). We observed that VDZ-responders had significantly smaller lymphoid aggregate size after treatment as compared to VDZ non-responders (Figure 6C, right), while pre-VDZ lymphoid aggregate size differences did not predict response to therapy (Figure 6C, left). In contrast to VDZ-treatment responders, we found no significant differences in lymphoid aggregate size among TNFi responders and non-responders after therapy (Figure 6D). These data suggest that the impact on lymphoid aggregate size was VDZ-specific and that the size loss was not dependent on treating tissue inflammation.

**Figure 6.**
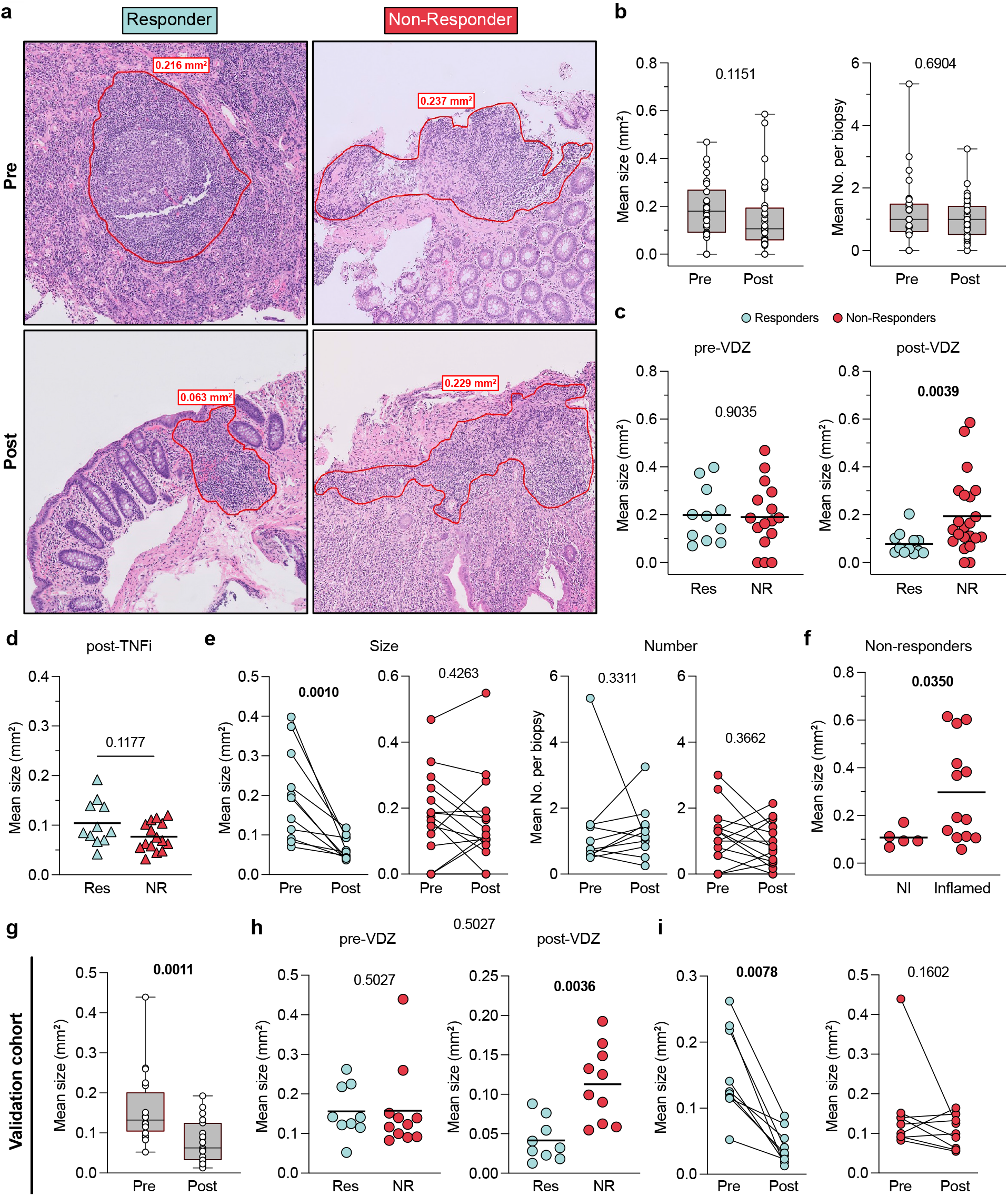
Response to vedolizumab is associated with loss of lymphoid aggregate size. **a**, Representative H&E staining of colonic biopsies from a Responder and a Non-Responder UC patient taken before and after VDZ therapy. Lymphoid aggregate (LA) border indicated with red line. **b**, Mean LA size and number per biopsy from UC samples taken before and after VDZ therapy. Data shown as Box-and-whisker plots. **c**, Comparison of LA size between VDZ Responders and Non-Responders in samples taken pre- and post-VDZ therapy. **d**, Mean LA size from UC patients after treatment with TNFi. **e**, Mean LA size (left) and mean LA number per biopsy (right) from paired samples taken pre- and post-VDZ therapy, from VDZ Responders and Non-Responders. **f**, Mean LA size from inflamed and non-inflamed tissue areas of VDZ Non-Responder UC patients after VDZ therapy. **g-i**, LA size analysis on an independent validation cohort. **g**, Mean LA size from UC patient samples taken before and after VDZ therapy. **h**, Comparison of LA size between VDZ Responders and Non-Responders in samples taken pre- and post-VDZ therapy. **i**, Mean LA size from paired samples taken pre- and post-VDZ therapy from VDZ Responders and Non-Responders. Paired non-parametric analysis were done using Wilcoxon test. Unpaired analyses were done using Mann-Whitney test. *p* values are indicated. Box plots represent median and quartiles of measurements, and whiskers represent minimum and maximum values.

Next, we studied longitudinal changes in lymphoid aggregate size, comparing colonic tissues pre- and post-VDZ in the same individuals. We observed a significant post-treatment reduction in lymphoid aggregate size in VDZ-responders (Figure 6E), remarkably, the size reduction was present in all responding patients. In contrast, there was no significant post-treatment change in lymphoid aggregate size in VDZ non-responders (Figure 6E). No significant change was observed in the number of lymphoid aggregates post-treatment in both VDZ responders and non-responders (Figure 6E). These findings suggest a strong association between lymphoid aggregate attrition and therapeutic response to VDZ.

We then focused on understanding if lymphoid aggregate size was related to local inflammation in VDZ-non responders. On comparing lymphoid aggregate size between inflamed and non-inflamed sites within the same patient, we observed that lymphoid aggregates were significantly larger in the inflamed areas compared to the uninflamed areas (Figure 6F). These observations suggest an association between ongoing local inflammation and lymphoid aggregate size.

To corroborate our findings in a geographically distinct population, we investigated the effect of VDZ on lymphoid aggregate size on an independent validation cohort of UC patients treated with VDZ (n=21), where colonic biopsies were obtained (COHORT 3 characteristics in Table S3). When we analyzed all patients, we observed a significant reduction in mean lymphoid aggregate size post-VDZ (Figure 6G). Further, while no difference was observed between responders and non-responders before treatment, we found that mean lymphoid aggregate size was significantly reduced in VDZ-responders as compared to non-responders post-treatment (Figure 6H). Finally, on longitudinal analysis of individual patients, biopsies from all VDZ-responders demonstrated a reduction in lymphoid aggregate size, in contrast to VDZ non-responders where no significant change in lymphoid aggregate size was seen post-treatment (Figure 6I). These observations from a geographically distinct validation cohort serve to corroborate our findings and suggest that the effect of VDZ on lymphoid aggregate size and its relation to therapeutic response could be generalizable to other populations.

## Discussion

Targeting the α4β7-MAdCAM-1 axis is a major component of IBD-therapeutics. VDZ, the prototypical anti-α4β7 antibody is now recommended for the induction and maintenance of remission in patients with moderate-to severe IBD^26^ and has rapidly emerged as a frontline biologic medication for patients with UC. The efficacy^27, 28^, unique safety profile^29^ and overall therapeutic success of VDZ has spurred interest in the development of several novel α4β7 antagonists^30^. This report details a major impact of VDZ on the GALT that relates to its therapeutic effect in two independent cohorts of patients with UC. VDZ binds to specific populations of circulating lymphocytes that include a subset of memory CD4^+^ T cells, naïve B cells, a subset of memory B cells, naïve CD4^+^ T cells, naïve and memory CD8^+^ T cells, NK cells, eosinophils and basophils^31^ in a pattern that is consistent with the expression profile of α4β7 among circulating mononuclear cells^3^. The therapeutic effect of VDZ was considered to be due to curtailed migration of effector T cells into the inflamed intestinal lamina propria^32, 33^. However, data on alterations in circulating and intestinal T cell frequencies post-VDZ are discordant. While Zeissig et al did not observe major differences in the absolute number, phenotype or TCR repertoire of mucosal and peripheral CD4, CD8, and central and effector memory T cells between VDZ-responders and non-responders^9^, subsequent studies by Coletta et al, noted a significantly reduced frequency of CD4 ^+^ CD127 ^+^ CCR6 ^+^ T cells and CD4^+^CD127^+^CCR6^+^CXCR3^+^ T cells (putative T^H17^ and T^H1^/T^H17^ cells respectively) in the lamina propria of VDZ-responders compared to non-responders^34^, while Veny et al observed a significant decrease in total T cells, CD4^+^ T cells and CD8^+^ T cells in the GI tract post VDZ, but detected no differences in T cell frequencies between treatment responders and non-responders^35^.

Taking an unbiased approach, we aimed to examine immune effector and inductive sites in VDZ-treated UC patients. While we did not discern significant changes in total or memory T cells and B cells, a significant decrease in naïve T cells and naïve B cells was evident in our data. Notably, a majority of intestine-resident T cells have a memory phenotype^36^, while naïve T cells and naïve B cells are enriched within GALT, including lymphoid aggregates and PP^37^. This prompted us to hypothesize that VDZ targets GALT as part of its therapeutic effect. Multiple lines of evidence support this hypothesis; first, α4β7 mediates the recruitment of naive T and B cells to gut-associated secondary lymphoid tissues^8, 38^. Second, VDZ significantly attenuates the humoral response to an oral cholera vaccine while preserving immune responses to a parenterally administered hepatitis B vaccine^39^. Third, when anti-α4β7 antibody was given to cynomolgus monkeys to assess for drug effect, while no significant change was noticed in the intestinal lamina propria, PPs were markedly atrophied^40^. Human data in the present report that are highly consistent with this hypothesis include a significant reduction in gut-homing plasmablasts in circulation and reduced frequencies of short-lived plasma cells in the lamina propria post-VDZ as both cell populations are induced within organized lymphoid tissues of the intestines^7^.

We tested our hypothesis in murine studies using multiple orthogonal experimental approaches and observed a highly significant attenuation of lymphoid structures following anti-α4β7 antibody administration, which was unequivocally associated with impaired entry of follicular naïve B cells and naïve T cells. Our data does not exclude some of the other suggested VDZ effects on intestinal and circulating immune cells that include reduced frequencies of mucosal DCs^41^ and a preferential inhibition of effector T cell migration over inhibition of T^REG^ migration^42^.

Lymphoid structures such as PP are anatomical sites of induction of antigen-specific immune responses^37^. Both human and murine data in our study suggest that attenuation of GALT by VDZ impacts humoral responses. While human studies showed a highly significant reduction in gut-homing plasmablasts in circulation post-VDZ, in murine experiments, we observed a significant decrease in OVA-specific IgA responses in OVA-gavaged mice that were treated with anti-α4β7-antibody compared to control mice. Further experiments on intestinal antigen-specific immune responses in human volunteers were outside the scope of the present report and will be planned in the future.

Finally, to define the clinical correlations of VDZ-associated GALT attenuation, we studied two additional patient cohorts-an internal Mount Sinai Cohort and an international cohort of well characterized VDZ-treated UC patients from Belgium. In both cohorts, a pathologist, blinded to the clinical details, scored lymphoid aggregate size in pre- and post-VDZ biopsies and determined that the reduction of lymphoid aggregate size correlated with response to VDZ. Lymphoid aggregate attenuation is likely VDZ-specific as we did not observe this in TNFi-treated humans or in murine studies with anti-TNF antibodies. Furthermore, although a decrease in lymphoid aggregate size was associated with resolution of inflammation, the VDZ effect on lymphoid structures is likely independent of inflammation as we noted a significant reduction in lymphoid aggregates not only in the colon but also in the (uninflamed) ileum of patients with UC. Moreover, it is unlikely that the size loss in lymphoid aggregates was due to the resolution of neighboring inflammation, considering that there was no difference in aggregate size between TNFi-responders and non-responders. Therefore, although we cannot indisputably infer causality between lymphoid aggregate size and tissue inflammation, our data does provide evidence supporting a model where attrition of lymphoid aggregates would contribute to limiting inflammation in the surrounding tissue.

In conclusion, we have identified the targeting of GALT as a novel mechanism of action of the anti-α4β7 antibody VDZ. By blocking the α4β7-MAdCAM-1 axis, VDZ prevents the entry of naïve B and T cells to GALT, resulting in loss of size and cellularity of these structures. GALT size loss is specific to VDZ and is associated with treatment response. Targeting of immune inductive sites represents a novel therapeutic paradigm in IBD care.

## Materials and Methods

### Study design

In this study, we aimed at characterizing the impact of anti-α4β7 therapy on the mucosal and circulating immune system of ulcerative colitis patients and to identify correlates of therapeutic response. We profiled immune cell changes in three distinct cohorts of ulcerative colitis (UC) patients, including one geographically distant patient cohort, where we obtained samples pre- and post-treatment. Patients were prospectively enrolled in the study, and tissue and blood samples were obtained, when possible, before and after vedolizumab therapy. As controls, we used samples taken from TNFi-treated and untreated UC patients. Histologic inflammation in evaluated by pathologists who were blinded to the patients’ treatment status (pre vs. post, type of medication, or responder vs. non-responder). Use of human samples was done in accordance with guidelines of the Mount Sinai Institutional Review Board. Furthermore, we studied mouse models of anti-α4β7 antibody treatment to get additional mechanistic insights. For this purpose, we used C57BL/6 and KikGR mice.

### Patients’ selection and clinical endpoints

For COHORT 1-2, patients were enrolled from the Inflammatory Bowel Disease (IBD) Center, the gastroenterology department and the digestive endoscopy unit at Mount Sinai Hospital, in accordance with local ethical guidelines. Informed consent was obtained from all participants. The study protocol was approved by the Mount Sinai institutional review board. Clinical characteristics are detailed in Tables S1-S2.

COHORT 1 (Mount Sinai): Patients with ulcerative colitis who started VDZ were prospectively enrolled (clinical characteristics in Table S1). In a subset of 43 patients for whom we performed FC characterization of PBMCs pre- and post-VDZ, blood was collected at week 0 and week 14 of treatment, just before the antibody infusion. In a subset of 12 patients, we collected ileal and colonic biopsies pre- and post-treatment during routine clinical care ileo-colonoscopies. All the pre-VDZ biopsies were collected less than one month before initiation of treatment. The remaining analysis on intestinal biopsies was performed in a cross-sectional manner at a single time point on patients receiving either no biologic (Untreated), TNFi or VDZ.

COHORT 3 (Belgium): This study was carried out at the University Hospitals Leuven (Leuven, Belgium). All included patients had given written consent to participate in the Institutional Review Board approved IBD Biobank of University Hospitals Leuven, Belgium (B322201213950/S53684).

### PBMCs isolation

Blood was drawn in in EDTA tubes and processed within 3 hours after collection. Blood was diluted with PBS (Gibco) and overlaid on lymphocyte separation medium (MP Biomedical). After centrifugation, PBMCs were collected and washed 2 times with PBS.

### Intestinal lamina propria cells isolation

During the colonoscopy, up to 30 biopsies from terminal ileum and left colon were collected, when possible, with forceps into ice-cold RPMI. Biopsies were processed within 3 hours of collection. Epithelial layer was removed by incubation (20 min, 37°C) in 10 ml HBSS (Ca-Mg-) containing EDTA (0.5M, pH 8, Invitrogen) and HEPES (1M, Lonza). Biopsies were vortexed, washed with HBSS and digested for 40 minutes at 37°C in a rotating incubator (215 rpm), in RPMI with 0.005 mg of collagenase (Sigma-Aldrich) and DNAse I (Sigma-Aldrich). Biopsies were physically disrupted using a 25-gauge needle, filtered through 100-µm and 40-µm cell strainers.

### Flow cytometry

The single-cell suspension was incubated with PBS containing the antibody cocktail for 20 minutes at 4°C in the dark. Cells were washed with PBS twice and fixed using Cytofix/Cytoperm buffer (BD). Cells were washed 2 times with Perm/wash buffer (BD). Cells were ultimately collected in FACS buffer (PBS w/o Ca2^+^ Mg2^+^ with 5mM EDTA) before acquisition. Samples were acquired using LSR Fortessa (BD Bioscience), and data was analyzed using FlowJo v10 (FlowJo). Analyses were all performed after the exclusion of dead cells and doublets. The list of used antibodies is displayed in Table S4.

### Histology

Hematoxylin & eosin-stained sections from colonic biopsies from COHORT 2 (Mount Sinai, Table S2) were scanned and viewed on a web-based platform (Philips IntelliSite Pathology Solution v. 3.3; Philips Medical Systems Nederland B.V., The Netherlands) and the area of each lymphoid aggregate was determined with the closed freeform tool. For the validation cohort (COHORT 3, Belgium, Table S3), biopsy slides were scanned into images (NDP.view2 software v. 2.9.29; Hamamatsu Photonics K.K., Japan) and lymphoid aggregate area was similarly determined using the freehand region tool. All lymphoid aggregates present on each slide were identified by virtue of their histomorphologic characteristics by experienced gastrointestinal pathologists (ADP and JW) and their individual areas were recorded. The degree of histologic inflammation in each biopsy was graded as previously described(43) by experienced gastrointestinal pathologists (ADP and JDP) who were blinded to the patients’ treatment status (pre vs. post, type of medication, or responder vs. non-responder).

### Mice

Animal care and experimentation were consistent with the US National Institutes of Health guidelines and approved by the Institutional Animal Care and Use Committee (IACUC) of the Icahn School of Medicine at Mount Sinai or IACUM at Washington University School of Medicine. C57BL/6 mice (B6) were purchased from Taconic Farms or bred and maintained in the Icahn School of Medicine facilities at Mount Sinai. KikGR mice were bred and maintained in the mouse facilities of Washington University. All animals were housed under specific pathogen-free (SPF) conditions and sacrificed at the indicated time points.

### Mouse injection and immunization

Six to eight-week-old mice were intra-peritoneally injected with anti-a4b7 IgG antibody (250µg in 250µl PBS) (Clone DATK32, Biolegend), anti-TNF IgG antibody (Clone MP6-XT22, Biolegend), isotype control IgG (RTK2758, Biolegend), or an equal amount of PBS.

For ovalbumin (OVA) immunization, mice were sensitized with an intraperitoneal injection of 500µg OVA (Sigma Aldrich) and 0.1µg cholera toxin (CT) (Sigma Aldrich) on day 0, followed by oral administration of 10mg OVA and 10µg CT on day 7 and 14. On day 21. Stool and blood were collected to measure OVA-specific IgA and OVA-specific IgG, respectively.

### Isolation of mouse Peyer’s patch cells and flow cytometry

Peyer’s patches (PPs) were individually collected and measured their weight. Each PP was incubated using collagenase D (Sigma-Aldrich, 400IU/ml) for 30 min and mechanically homogenized for single-cell suspension. After Fc blocking using CD16/32 antibody (2.4 G2, BioXcell), cells were stained with fluorescent conjugated antibodies (Table S5) for flow cytometry. For calculating the total cell number from each PP, 5 µL of AccuCheck Counting Beads (Invitrogen) were added to each sample before acquiring the data. Data was acquired using LSR Fortessa (BD Bioscience) and analyzed using FlowJo v10 (FlowJo).

### Mouse Immunofluorescence

Tissues were fixed in 4% paraformaldehyde in PBS overnight at 4°C, washed three times for 30 min in PBS, then moved to 30% sucrose in PB overnight. Tissues were flash frozen in Tissue-Tek Cryomold (VWR) the next day, and 7-mm sections were cut and then dried for 1 hour before staining. Sections were rehydrated in PBS with 1% bovine serum albumin (BSA) for 10 min and then stained overnight at 4C and stained for subsequent steps at room temperature for two hours, all in PBS with 1% BSA, 2% mouse serum, 2% rat serum and 2% donkey serum. Sections were stained with primary antibodies: Goat anti-mouse IgD (goat polyclonal GAM/IgD, FC/7S, Cedarlane Labs), biotin anti mouse GL7 (Biolegend). Sections were then stained with the secondary antibodies: Cy3-Streptavidin (Jackson Immunoresearch), AF488 anti-goat (Jackson Immunoresearch), and 40,6-diamidino-2-phenylindole (DAPI, ThermoFisher). ImageJ software was used for area quantification.

### Hashtag Antibody Staining

Cells isolated from Peyer’s patches were counted using the Nexcelom Cellometer Auto 2000. An aliquot of 1,000,000 cells from each sample was centrifuged at 400 RCF for 10 minutes (4°C). The supernatant was discarded and the pellets were resuspended in 100ul of 10x Genomics Cell Multiplexing Oligo (CMO) and incubated at RT for 5 minutes. Stained cells were washed three times in 1mL of cold wash buffer (PBS + 1% BSA) by 4°C centrifugation at 400 RCF to remove unbound CMOs. Washed cells were resuspended in 200ul resuspension buffer (PBS + 0.05% BSA) and counted. Stained samples were pooled in equal cell numbers and centrifuged at 400 RCF for 5 minutes (4°C). Supernatant was removed and pellet was resuspended in 100ul of resuspension buffer.

### Single-cell RNA sequencing

Sample pool was counted and loaded on one lane of the 10x Genomics NextGem 3’v3.1 assay as per the manufacturer’s protocol with a targeted cell recovery of 24,000 cells per lane. Gene expression and CMO libraries were made as per the 10x Genomics demonstrated protocol (https://cdn.10xgenomics.com/image/upload/v1666737555/support-documents/CG000388_ChromiumNextGEMSingleCell3-v3.1_CellMultiplexing_RevC.pdf).

All libraries were quantified via Agilent 4200 Tapestation High Sensitivity D5000 ScreenTape Assay (Agilent Cat# 5067-5592, 5067-5593) and KAPA library quantification kit (Roche Cat# 0796014001.) Gene expression libraries were sequenced at a targeted depth of 25,000 reads per cells. CMO libraries were sequenced at a targeted read depth of 1,000 reads per cell. Libraries were sequenced on the Illumina NovaSeq S2 100 cycle kit with run parameters set to 26/10/10/90 (R1xi7xi5xR2).

### Processing single-cell RNA data

BCL files were demultiplexed into FASTQs using Cell Ranger v6.1.2. Alignment, filtering, barcode counting, and UMI counting were also performed using Cell Ranger v6.1.2. Samples were aligned to mouse reference mm10-2020-A.

Cells were filtered based on total number of UMI counts and mitochondria genes fraction. Specifically, only cells with total UMI counts greater than 500 and mitochondria gene fraction less than 10% were considered for the analysis. The analysis of single-cell RNA data including data normalization and batch effect correction was performed via the Seurat package(44). First, each data was normalized independently using function NormalizeData which divides feature counts for each cell by the total counts for that cell. Data was then natural-log transformed. Next, we found variable features using function FindVariableFeatures(44). This function first fits a line to the relationship of log(variance) and log(mean) using local polynomial regression; then, standardizes the feature values using the observed mean and expected variance (given by the fitted line). Feature variance was then calculated based on the standardized values. Finally, we selected features which were repeatedly variable across datasets for integration via function SelectIntegrationFeatures. We then identified anchors using the FindIntegrationAnchors() function, which takes a list of Seurat objects as input, and use these anchors to integrate different datasets together via the IntegrateData function(44). Dimensionality reduction to identify anchors was performed using reciprocal principal component analysis.

### Clustering single-cell RNA data

Clustering was performed via function Find.Cluster available in the Seurat package(44). This function identifies clusters of cells using a shared nearest neighbor algorithm. Once that unsupervised clustering was performed, we identified cluster-specific markers using function FindMarkers(44). Markers were ranked based on area under the receiver operating characteristic curve (AUC). Based on these cluster-specific markers, clusters were annotated into 15 cell-types.

### Differential expression between treated and untreated mice

Differential expression between treated and untreated mice within a particular cell-type was performed using function FindMarkers(44). Markers were ranked based on area under the receiver operating characteristic curve (AUC) and only markers with an AUC greater than 0.7 were considered as differentially expressed between treated and untreated mice.

### Mouse stool preparation and ELISA

Murine stool (50 mg) was incubated in 500 µL of PBS at room temperature for 10 minutes, followed by vortexing for 5 minutes and bead-beating for 2 minutes. The sample supernatants were collected after centrifugation.

ELISA was performed using Mouse IgA ELISA Quantitation Set or Mouse IgG ELISA Quantitation Set (Bethyl Laboratories) according to the manufacturer’s protocol with minor modifications. MaxiSorp ELISA plates (Thermo Fisher Scientific) for total IgA measurement were pre-coated with 100 µL of diluted purified IgA antibody in 0.1M carbonate buffer (10 µl/mL). For OVA-specific IgA or IgG measurement, MaxiSorp ELISA plate were pre-coated with 5µg of OVA protein in 0.1M carbonate buffer. After overnight incubation at 4°C, plates were washed with PBS and incubated with PBS containing 1% BSA for 1 hour at room temperature. After washing, 100 µL of stool supernatant or serum were plated and incubated for 1 hour. Plates were washed again and 100 µL of anti-mouse IgA-HRP or anti-mouse IgG HRP (1:40 000 dilutions) was added before a 1-hour of incubation. TMB substrate (BD Pharmingen) was added to each well, followed by supplementation of 100 µL of 1M H^2^SO4 as a stop solution. Absorbance (450 nm) was measured using an ELISA reader.

### In-situ photoconversion

Peyer’s patch (PP) photoconversion labeling was performed as previously reported(25). In brief, mice were anesthetized with 2% isoflurane during the procedure. An abdominal wall incision exposed the PP. An opaque, non-reflective material shielded the mouse from light exposure except at the desired site. Photoconversion of the three most distal PP was performed using a low-power 80mW LED-Violet laser (Laserland) in 3 cycles of 10 seconds powered on followed by 30 seconds powered off, keeping the light source approximately 2 cm away from the tissue. Next, mice were sutured and allowed to recover from anesthesia on a heating pad until responsive. Just prior to surgery, mice were given long-acting buprenorphine for analgesia. Two hours before the photoconversion, mice were treated intraperitoneally with 100 ug of α4β7 (DATK32, Biolegend; cat# 120602) or Rat IgG2a isotype control (RTK2758, Biolegend; cat# 400502) antibodies. PPs were then harvested 20 hours after photoconversion. KikRED^+^ cells (in situ labeled cells) and KikGREEN^+^ cells were quantified in all small intestinal PP by flow cytometry.

### Statistical analyses

Data is shown as individual points and mean, paired before-after plots or as box-and-whisker plots. Unpaired analysis was done using Mann-Whitney test. For multiple comparisons, non-parametric analysis was done using Kruskal-Wallis test and Dunn’s multiple comparison test. Paired non-parametric analysis was done using Wilcoxon test. The p values are indicated. For single-cell RNAseq data analysis, see specific section.

## Supplementary materials

Supplementary Figures S1 – S5 Supplementary Tables S1 – S5

## Acknowledgements

We thank the patients who participated in this study. We would like to thank the Human Immune Monitoring Center at Mount Sinai for carrying out single-cell RNA-seq experiments.

## Author contributions

Conceptualization: PCH, MU, AS, JFC, SM. Investigation: PCH, MU, AS, RSC, BV, AL, FR, AD, AW, DD, MT, DJ, AK, JDP, GC, TD, ADP. Visualization: PCH, MU, AS, RSC. Funding acquisition: SM, JFC, AR, ADP. Supervision: SM, JFC, AR, ADP. Writing: PCH, MU, SM.

## Data materials availability

All data needed to support the conclusions of this work are included in the main text and supplementary materials.

## Supplementary materials

**Figure S1.**
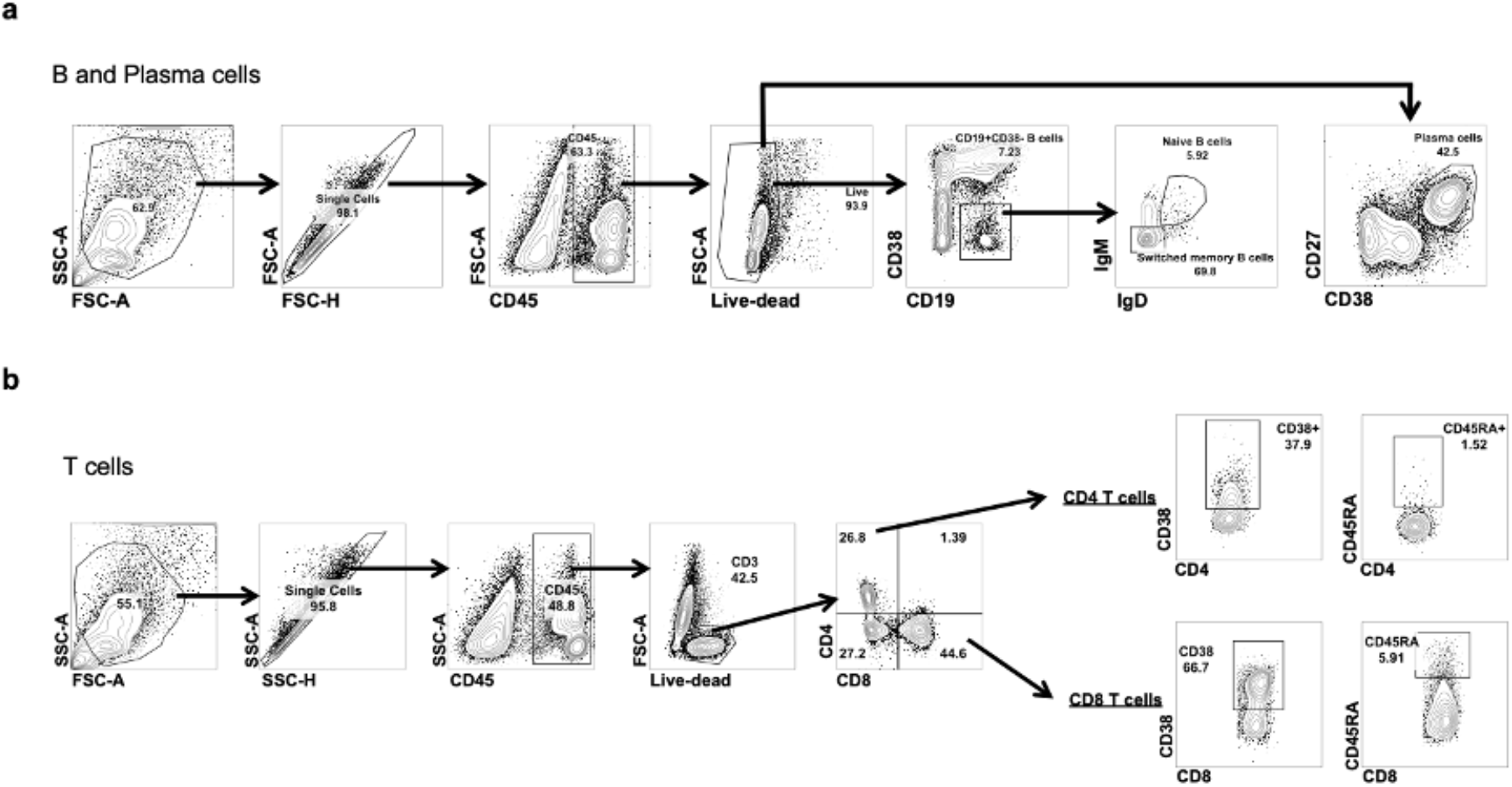
Gating strategy for human intestinal cells. **a-b**, Representative flow cytometry (FC) plots showing gating strategy used to profile intestinal B cells and plasma cells (**a**), as well as T cells (**b**).

**Figure S2.**
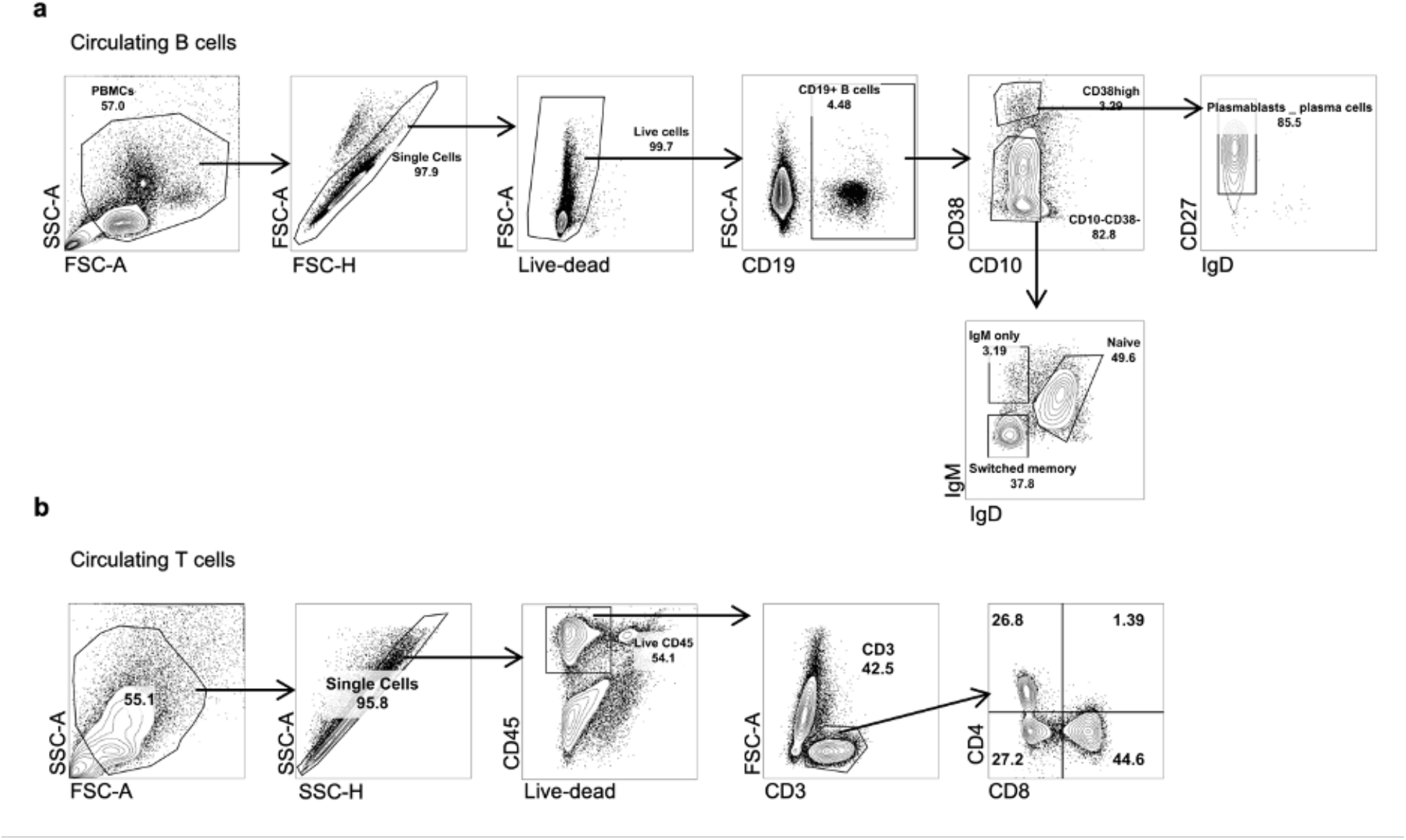
Gating strategy for human circulating cells. **a-b**, Representative flow cytometry (FC) plots showing gating strategy used to profile circulating B cells and plasma cells (**a**), as well as T cells (**b**).

**Figure S3.**
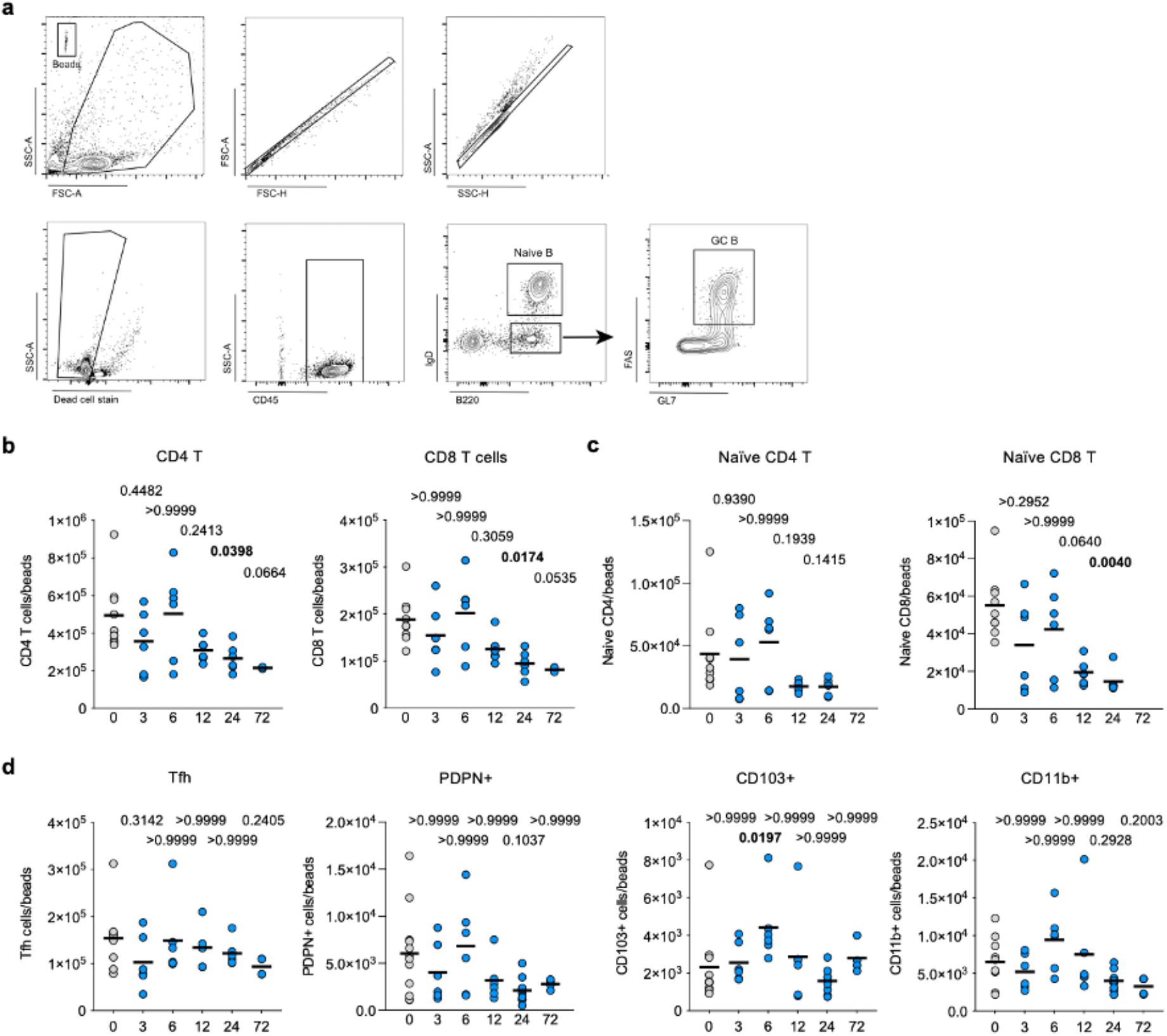
Kinetic profiling of mouse PPs after anti-α4β7 administration. **a**, Representative flow cytometry (FC) plots showing gating strategy used to profile mouse B cells. **b**, Frequency of CD4 T cells and CD8 T cells from isolated PPs of untreated mice (grey) and at 3, 6, 12, 24 and 72 hours after anti-α4β7 administration. **c**, Frequency of Naïve CD4 and CD8 T cells after anti-α4β7 administration. **d**, Frequency of T follicular helper, PDPN+, CD103+ and CD11b+ cells after anti-α4β7 administration. Data shown as individual values and mean. Unpaired non-parametric analysis was done using Kruskal-Wallis test and Dunn’s multiple comparisons test. The *p* values are indicated.

**Figure S4.**
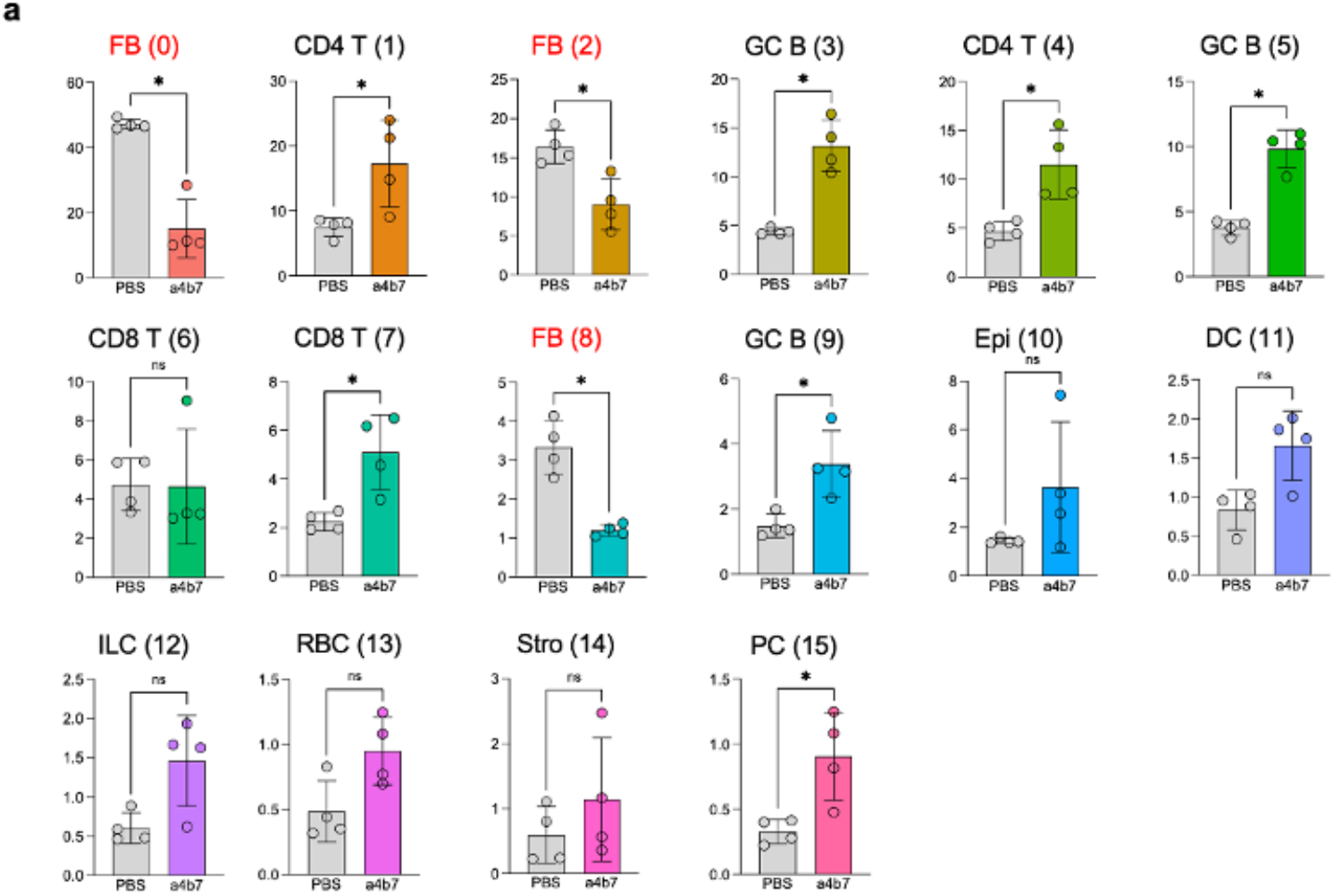
Frequency of single-cell RNAseq clusters. **a**, Bar plots for the frequency of cell clusters among total cells from Figure 4. Data shown as individual values and mean. Unpaired analysis was done using Mann Whitney test.

**Figure S5.**
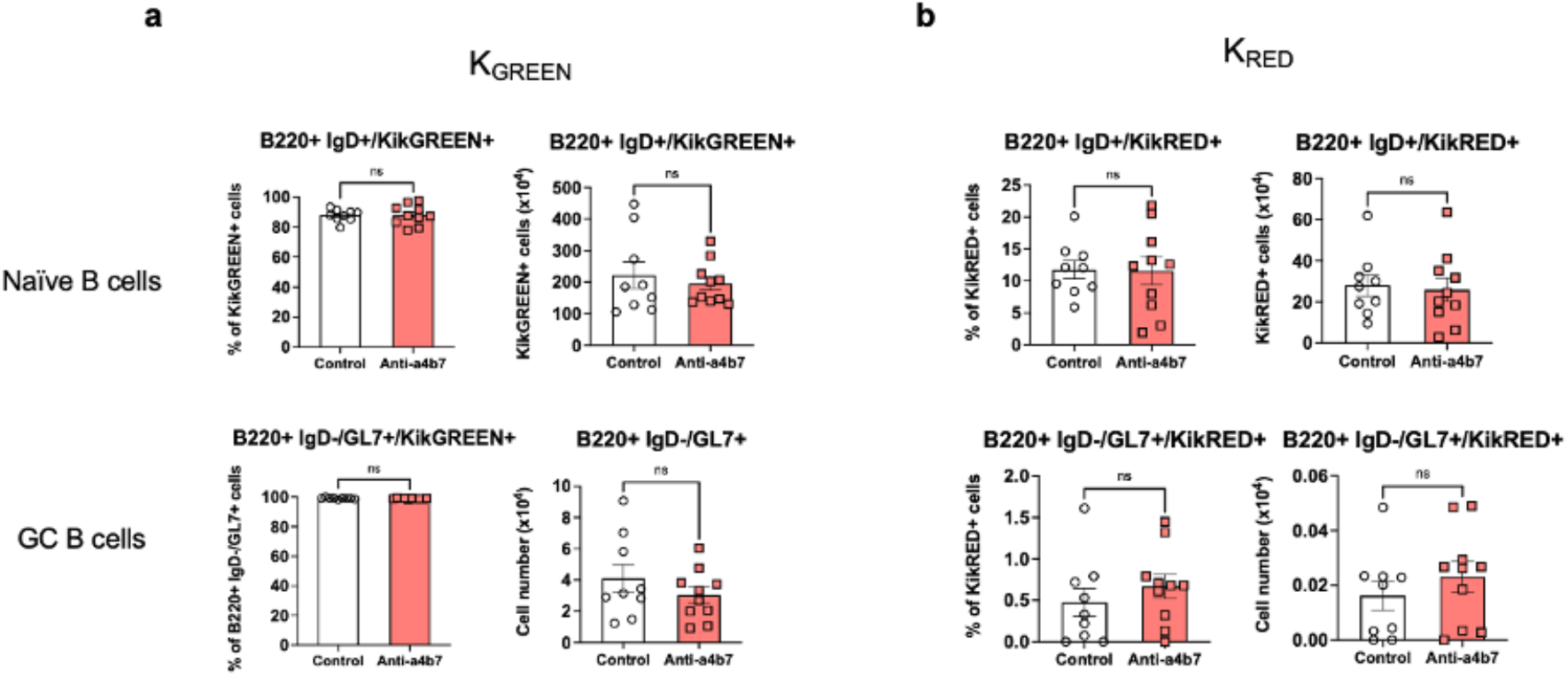
B cell monitoring in mesenteric lymph nodes after photoconversion. Cell frequency and absolute cell number of K^GREEN^ (**a**) and K^RED^ (**b**) Naïve B cells and GC B cells from non-photoconverted mesenteric lymph nodes. Mice were treated with anti-α4β7 or Isotype control as in Figure 5. Data shown as individual values, mean and SD. Unpaired analysis was done using Mann Whitney test.

**Table S1.** COHORT 1 (Mount Sinai) used for flow cytometry studies.

|  | **Vedolizumab**<br>**treated** | **TNFi treated** | **Untreated** | **p-value** |
| --- | --- | --- | --- | --- |
| Number of patients | 56 | 10 | 17 | NA |
| Age (mean ± SD) | 38.39 ± 13.43 | 29.94 ± 9.72 | 40.92 ± 19.1 | 0.15 |
| Sex, n M / F | 29 / 27 | 15 / 7 | 13 / 4 | 0.01 |
| Disease activity/extent*, n |  |  |  |  |
| Not active | 3 | 2 | 3 |  |
| E1 | 5 | 1 | 5 |  |
| E2 | 22 | 4 | 6 |  |
| E3 | 26 | 3 | 3 | 0.1 |

\*extent of active disease at time of endoscopy
SD = Standard deviation. Age is at the time of first biopsy analyzed. For continuous variables used One-way ANOVA, for categorical variables used Chi-square test for trend.

**Table S2.**
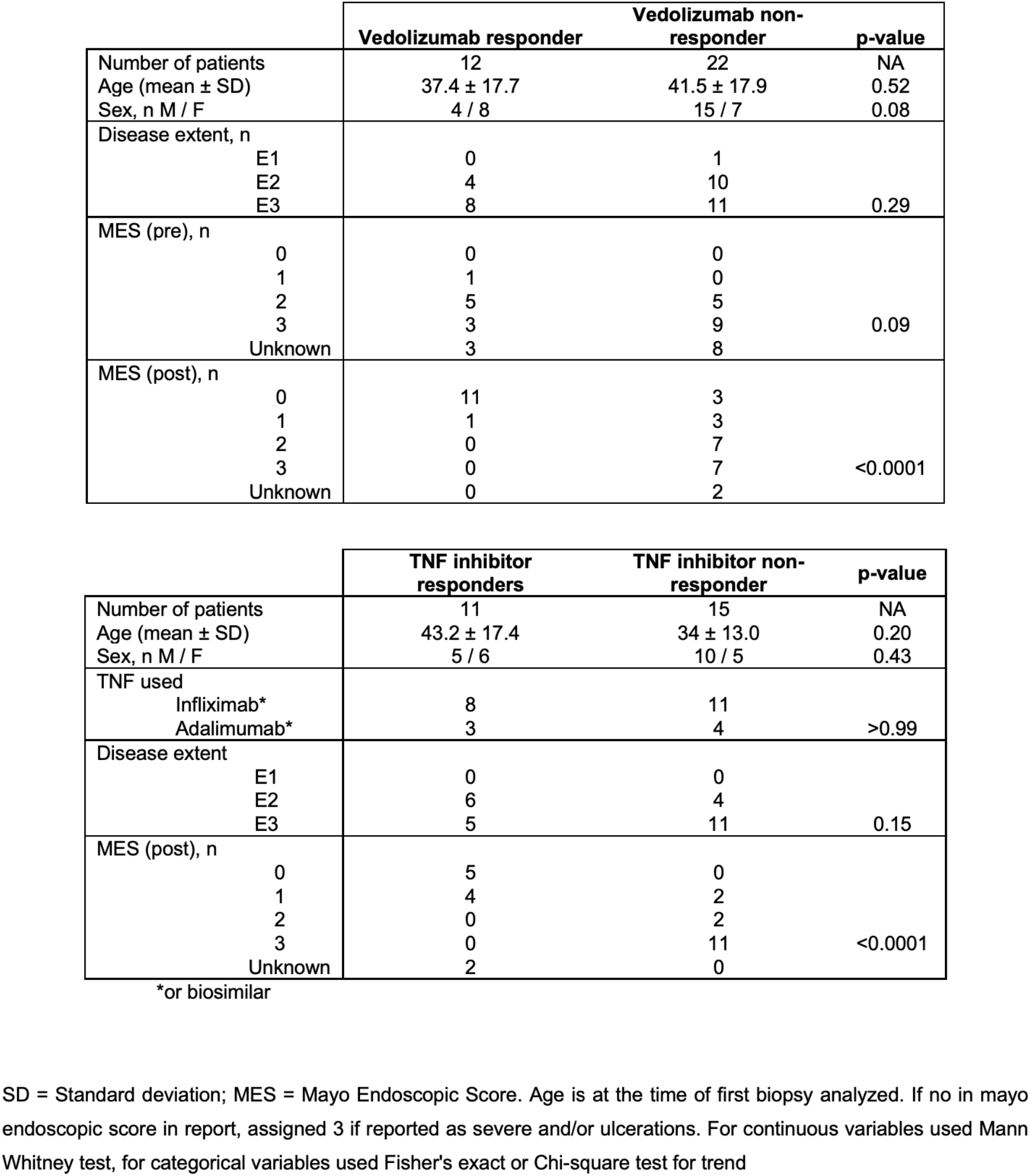
COHORT 2 (Mount Sinai) used for histology studies.

**Table S3.** COHORT 3 (Validation cohort, Belgium) used for histology studies.

|  | Vedolizumab responder | Vedolizumab non-responder | p-value |
| --- | --- | --- | --- |
| Number of patients | 9 | 12 | NA |
| Age (mean ± SD) | 50.8 ± 18.36 | 43.27 ± 10.8 | 0.24 |
| Sex, n M / F | 5 / 4 | 7 / 5 | >0.9999 |
| Disease extent, n |  |  |  |
| E1 | 3 | 2 |  |
| E2 | 5 | 5 |  |
| E3 | 1 | 5 | 0.13 |
| MES (pre), n |  |  |  |
| 0 | 0 | 0 |  |
| 1 | 0 | 0 |  |
| 2 | 1 | 6 |  |
| 3 | 8 | 6 | 0.06 |
| Unknown | 0 | 0 |  |
| MES (post), n |  |  |  |
| 0 | 4 | 3 |  |
| 1 | 3 | 2 |  |
| 2 | 2 | 1 |  |
| 3 | 0 | 6 | 0.04 |
| Unknown | 0 | 0 |  |

SD = Standard deviation; MES = Mayo Endoscopic Score. Age is at the time of first biopsy analyzed. If no in mayo endoscopic score in report, assigned 3 if reported as severe and/or ulcerations. For continuous variables used Mann Whitney test, for categorical variables used Fisher's exact or Chi-square test for trend.

**Table S4.**
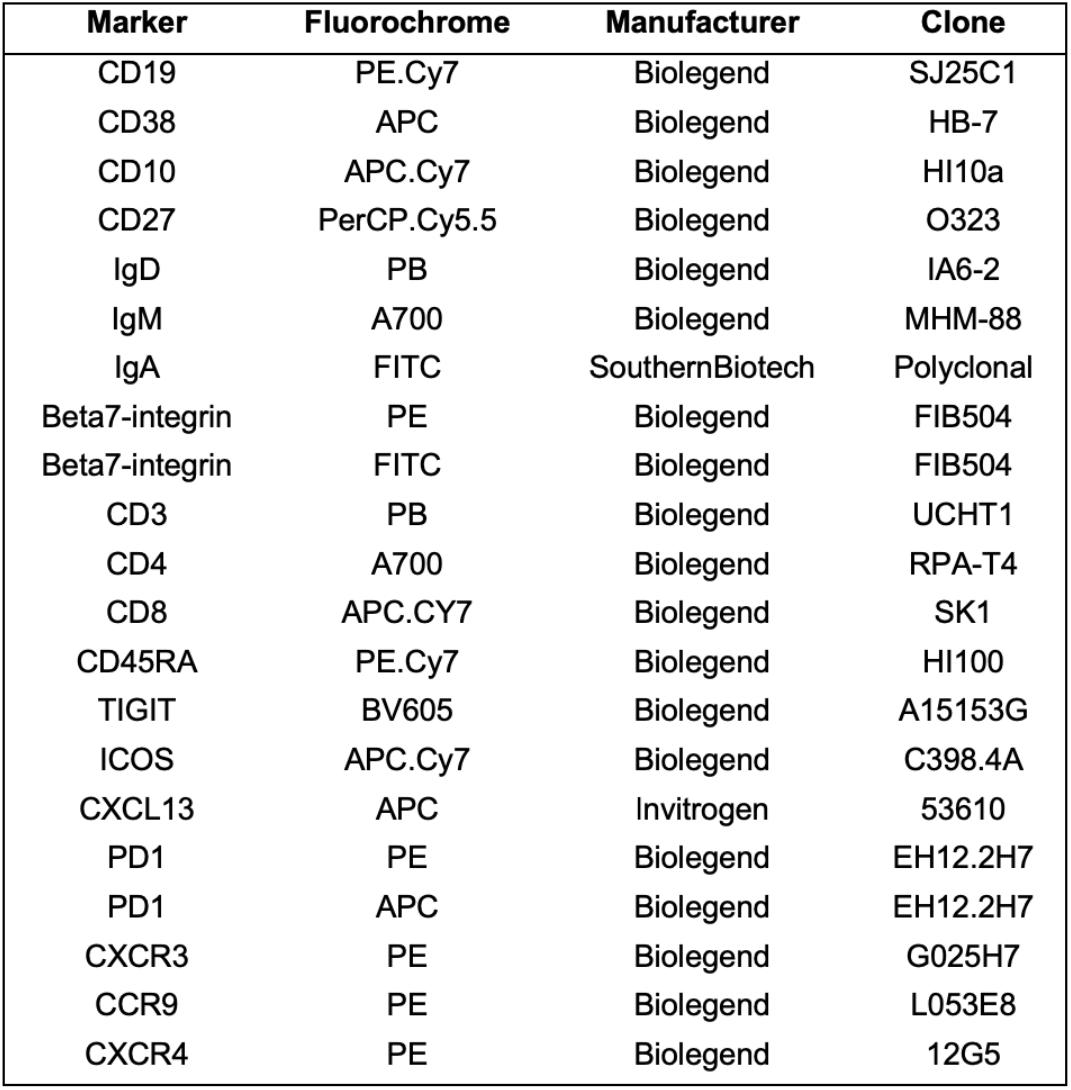
Antibodies used for Flow cytometry of human samples.

**Table S5.**
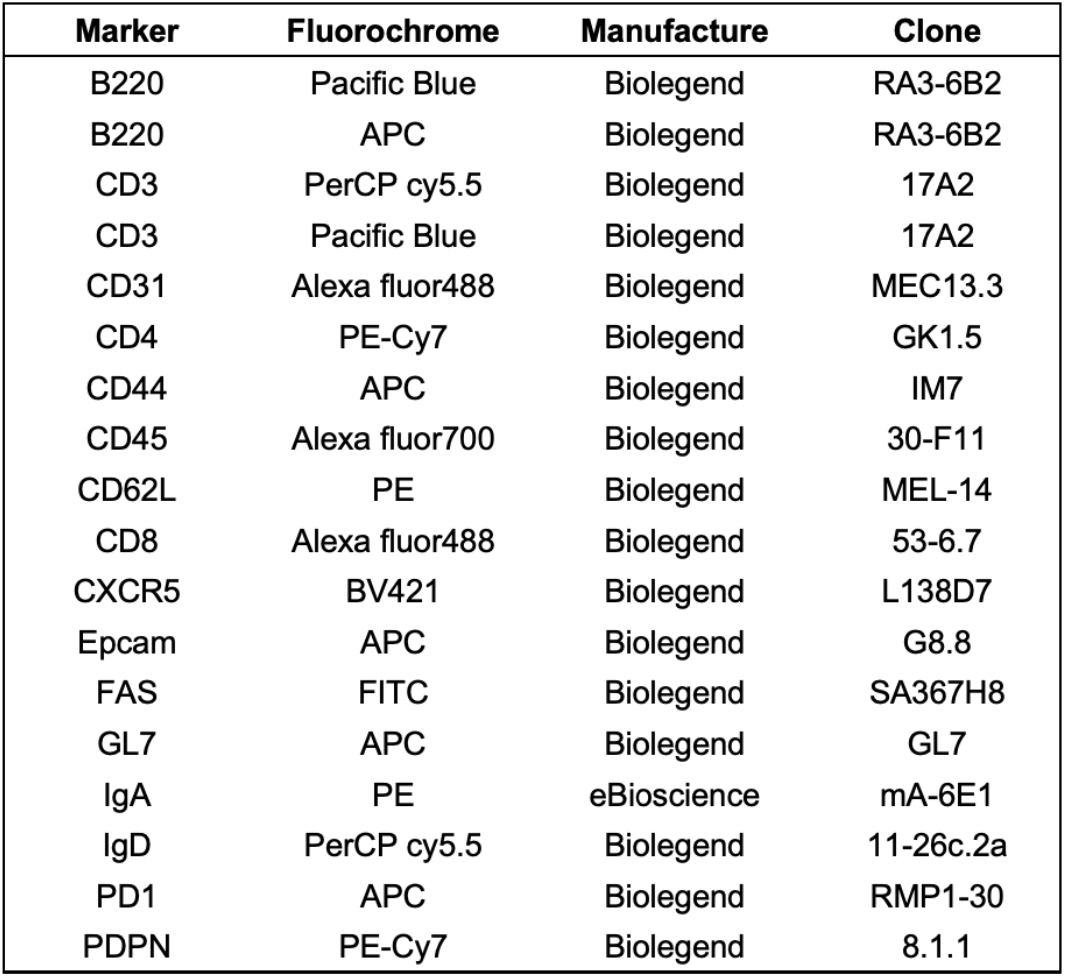
Antibodies used for Flow cytometry of murine samples.

| Marker | Fluorochrome | Manufacture | Clone |
| --- | --- | --- | --- |
| B220 | Pacific Blue | Biolegend | RA3-6B2 |
| B220 | APC | Biolegend | RA3-6B2 |
| CD3 | PerCP cy5.5 | Biolegend | 17A2 |
| CD3 | Pacific Blue | Biolegend | 17A2 |
| CD31 | Alexa fluor488 | Biolegend | MEC13.3 |
| CD4 | PE-Cy7 | Biolegend | GK1.5 |
| CD44 | APC | Biolegend | IM7 |
| CD45 | Alexa fluor700 | Biolegend | 30-F11 |
| CD62L | PE | Biolegend | MEL-14 |
| CD8 | Alexa fluor488 | Biolegend | 53-6.7 |
| CXCR5 | BV421 | Biolegend | L138D7 |
| Epcam | APC | Biolegend | G8.8 |
| FAS | FITC | Biolegend | SA367H8 |
| GL7 | APC | Biolegend | GL7 |
| IgA | PE | eBioscience | mA-6E1 |
| IgD | PerCP cy5.5 | Biolegend | 11-26c.2a |
| PD1 | APC | Biolegend | RMP1-30 |
| PDPN | PE-Cy7 | Biolegend | 8.1.1 |

